# Regulatory fine-tuning and horizontal gene transfer stabilize mobile colistin resistance

**DOI:** 10.1101/2022.11.04.515217

**Authors:** Lois Ogunlana, Liam Shaw, Divjot Kaur, Pramod Jangir, Timothy Walsh, Stephan Uphoff, R.C. MacLean

**Affiliations:** Department of Biology, University of Oxford. 11a Mansfield Road, Oxford OX1 3SZ; Ineos Oxford Institute for Antimicrobial Research, University of Oxford, South Parks Road, Oxford OX1 3RE; Department of Biochemistry, University of Oxford, South Parks Road, Oxford OX1 3QU

## Abstract

Antibiotic resistance tends to carry fitness costs, making it difficult to understand how resistance can be stably maintained in pathogen populations over the long-term. Here, we investigate this problem in the context of *mcr-1*, a fitness-costly gene that confers resistance to the ‘last-resort’ antibiotic, colistin. Here we show that regulatory evolution has fine-tuned the expression of *mcr-1*, allowing *E. coli* to reduce the cost of *mcr-1* while simultaneously increasing colistin resistance. Conjugative plasmids have transferred low cost/high resistance *mcr-1* alleles across an incredible diversity of *E. coli* strains, further stabilizing *mcr-1* at the species level. Crucially, regulatory mutations were associated with increased *mcr-1* stability in pig farms following a ban on the use of colistin as a growth promoter that decreased colistin consumption by 90%. Our study shows how the rapid evolution and horizontal transmission of resistance genes can combine to stabilize resistance and reduce the efficiency of interventions aimed at reducing AMR by limiting antibiotic consumption.

## Introduction

Antibiotic resistance in pathogenic bacteria has emerged as a fundamental threat to human health, prosperity and food security (Murray et al. 2022; Aminov 2010; Levy and Bonnie 2004). In light of this threat, it is vital to develop a thorough understanding of the processes that drive the rise and fall of resistance in pathogen populations. Exposure to antibiotics generates selection for resistance, providing pressure for the spread of resistance genes in pathogen populations (Stanton et al. 2020; Gullberg et al. 2011; Aarestrup 2005; MacLean et al. 2010; Palmer and Kishony 2013; Andersson et al. 2020). However, a key challenge in this area is to understand how antibiotic resistance can stably persist in pathogen populations, including in the absence of antibiotic use (Andersson and Hughes 2010; MacLean and Vogwill 2015; Andersson et al. 2020).

Antibiotic resistance is often associated with fitness costs, such as reduced competitive ability and virulence, making it challenging to understand how resistance can persist in pathogen populations over the long term (Vogwill and Maclean 2015; Melnyk, Wong, and Kassen 2015; Andersson and Hughes 2010; Andersson and Levin 1999). For example, resistance may hinder the ability of pathogenic strains to transmit between people via environmental reservoirs where antibiotic concentrations are low, such as fomites or contaminated drinking water, and resistance often declines in patients following antibiotic treatment (Chung et al. 2022; Wheatley et al. 2021; B. G. Bell et al. 2014; Malhotra-Kumar et al. 2007). Given these costs, one of the simplest strategies to combat antibiotic resistance is to reduce antibiotic consumption via stewardship programs (Basra et al. 2018; G. Bell and MacLean 2018; Andersson et al. 2020; Andersson and Hughes 2010).

In contrast to this expectation, experimental evolution studies usually find that resistance stably persists in the absence of antibiotic exposure (Baltrus 2013; Andersson and Hughes 2010; Millan et al. 2014; Andersson and Levin 1999; Schrag, Perrot, and Levin 1997; Levin et al. 1997). Resistance is stably maintained in these experiments by compensatory mutations that offset the costs of resistance. The idea that resistance is stabilized by compensatory adaptation is deeply ingrained in evolutionary models of antibiotic resistance (Andersson and Levin 1999; Millan et al. 2014; Andersson 2003; zur Wiesch et al. 2011; MacLean and San Millan 2019; Andersson et al. 2020). However, direct examples of resistance being stabilized by compensatory adaptation in clinical pathogen populations are lacking (MacLean and Vogwill, 2015), with the notable exception of studies on *M. tuberculosis* (Comas et al. 2012; Gygli et al. 2021). Moreover, alternative mechanisms can stabilize resistance without compensatory adaptation. For example, resistance genes can persist as a result of selection for linked genes, such as biocide resistance genes, a phenomenon called co-selection (Andersson and Hughes 2010). Co-selection is particularly important when high-fitness pathogen strains acquire resistance genes. Many resistance genes are carried on mobile genetic elements, particularly conjugative plasmids. High rates of horizontal transfer of these elements can allow resistance to persist in pathogen populations, even if the elements themselves impose a small fitness cost (Stewart and Levin 1977; Lopatkin et al. 2017; Hall et al. 2016).

The aim of this study was to understand how colistin resistance is stabilized in *Escherichia coli*. The widespread use of colistin as an agricultural growth promoter drove the rapid spread of colistin resistant *E. coli* across one-health settings, including farms, humans and the environment (Y. Shen, Zhou, et al. 2018). Colistin is a WHO reserve antibiotic for the treatment of infections caused by multi-drug resistant gram-negative pathogens (Adekoya et al. 2021). The widespread presence of colistin resistance in pathogenic and commensal *E. coli* represents an important threat to the clinical utility of colistin.

Colistin resistance in *E. coli* has been driven by the acquisition of **m**obile **c**olistin **r**esistance (i.e., MCR) genes that are mainly carried on conjugative plasmids (Liu et al. 2016; Sun et al. 2018). Many *mcr* homologues have now been identified, but *mcr-1* remains the most prevalent and best-characterized colistin resistance gene (Sun et al. 2018; Shen et al. 2020). *mcr-1* initially spread as part of a composite IS*ApI*1 transposon that transferred between plasmids, which themselves transferred between strains of pathogenic and commensal *E*.*coli* and other enteric bacteria (R. Wang et al. 2018). *mcr-1* expression results in fundamental changes to the bacterial outer membrane, leading to extensive pleiotropic effects (Yang et al. 2017; Li et al. 2020). Unsurprisingly, these modifications are associated with large fitness costs. For example, expression of *mcr-1* can reduce the fitness of *E*.*coli* by as much as 30% (Yang et al. 2017). To put this in context, plasmids carrying mobile resistance genes typically have fitness costs on the order of 10% (Vogwill and MacLean, 2014)

The Chinese government responded to the spread of *mcr-1* by banning the use of colistin as a growth promoter in animal feed in April 2017, resulting in a 90% reduction in colistin consumption (Shen et al. 2020; Wang et al. 2020; Walsh and Wu 2016). Large-scale surveillance studies found that the prevalence of *mcr-1* declined following the colistin ban, but in many sectors the decline was much slower than would be expected given the significant fitness cost of colistin resistance previously been reported (Yang et al. 2017). For example, the prevalence of colistin-resistant *E. coli* declined in pigs from 34% pre-ban (2015-16) to 5.1% post-ban (2017-2018) (Wang et al. 2020). If we conservatively assume that *E. coli* has a generation time of approximately 12 hours (Barroso-Batista, Demengeot, and Gordo 2015; Myhrvold et al. 2015), this decline over a two-year period spanning >1,000 generations would suggest a fitness cost of <1%. This implies that *mcr-1* is much more stable than we would expect in natural populations than is implied by fitness costs measured *in vitro*.

The goal of this paper is to understand how *mcr-1* has been stabilized using a combination of genomics, competition assays and epidemiological data. Using this integrative approach, we first identify common polymorphisms in the regulatory region of *mcr-1* which existed before the colistin ban. We then demonstrate that these regulatory mutations reduce the cost of *mcr-1* carriage by fine tuning its expression. Unlike classical compensatory mutations, these regulatory mutations also confer increased colistin resistance. Importantly, conjugative plasmids have transferred *mcr-1* genes with fine-tuned expression across different strains of *E. coli*, providing evidence that horizontal gene transfer further contributes to stabilizing *mcr-1*. Finally, we show that regulatory mutations have stabilized *mcr-1* in pig-associated populations following the ban on the use of colistin as a growth promoter.

## Results

### Regulatory polymorphisms alleviate the cost of *mcr-1*

A comprehensive genomic analysis of MCR positive *E. coli* published in 2018 identified a polymorphism hotspot upstream of the *mcr-1* gene (R. Wang et al. 2018). Examination of this region revealed the presence of multiple polymorphisms in regions that are predicted to encode RNA polymerase binding sites (i.e. -10 and -35 boxes) and the Shine-Dalgarno (SD) site (Figure.1A, Table.S1). The presence of parallel nucleotide substitutions in this reported regulatory region of *mcr-1* suggests that these polymorphisms represent selectively advantageous mutations (Lieberman 2022). To directly test this hypothesis, we cloned *mcr-1* into pSEVA121 under the control of either the wild-type (WT) regulatory sequence (i.e., the consensus sequence) or one of eight known mutated ‘regulatory variant’ sequences (Figure.1A). These variants were associated with a single polymorphism within the -10 promoter or Shine-Dalgarno (SD) region apart from SDV2 which had two polymorphisms within the SD region. The pSEVA121 vector used in our experiments has a similar copy number (approximately 4-5/cell) to natural plasmids that carry *mcr-1* (typically 2-5 per cell).

**Figure 1:**
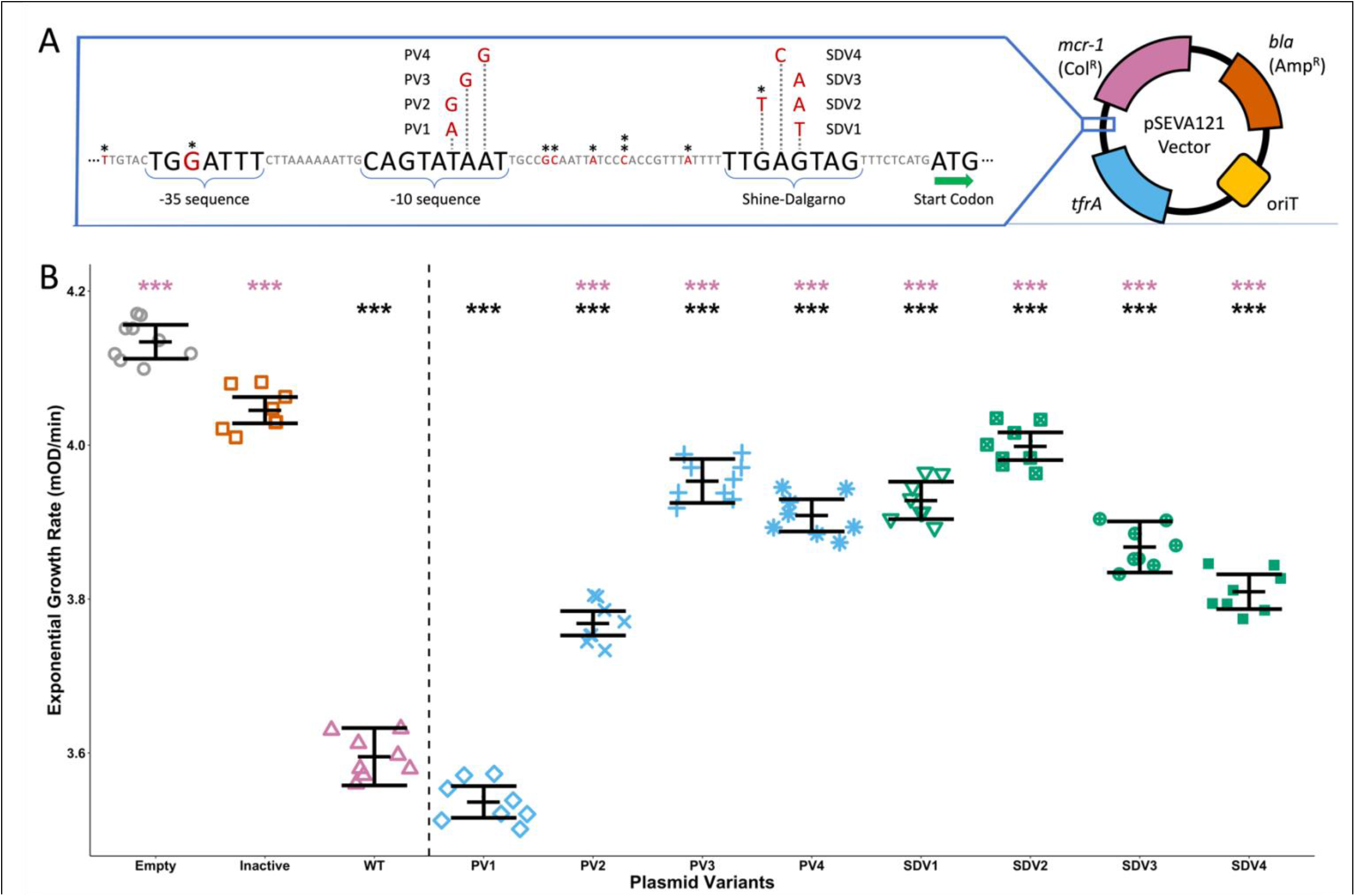
Construction and Fitness assessment of *mcr-1* regulatory variants. **A**) Schematic shows the region immediately upstream of the *mcr-1* start codon. Canonical regulatory regions are labelled, with polymorphisms shown in red. Polymorphisms found in each tested regulatory variant are named. Asterisks denote sites where SNPs were found. Regulatory variants and a WT regulatory sequence were cloned into the pSEVA121 vector. **B**) Exponential growth rates (mOD/min) in colistin-free media are shown for the WT regulatory sequence (WT, pink) and promoter (blue) and Shine-Dalgarno (green) variants. Empty (grey) inactivated *mcr-1* (orange) controls are included. The experiment was replicated over 8 different days. Each plotted point shows the average growth rate of 5 replicate cultures in a single run of the experiment. Significance of comparisons to empty (black) and WT (pink) controls are indicated (p-values: ***<0.001. Dunnett’s t-test adjusted for multiple comparisons, Error bars = standard error, n = 5).

The large fitness costs associated with *mcr-1* are likely to generate strong selection for mutations that reduce the cost of *mcr-1*. To test this idea, we first measured the growth rate of strains carrying regulatory variants and wild-type strains in colistin-free culture medium, which is a common method to assess the fitness costs of resistance (Vogwill and Maclean 2015). As a control, we also measured the fitness effect of an inactivated variant which carried a wild-type regulatory sequence and a mutated *mcr-1* active site (T285A) (Stojanoski et al. 2016; Ma et al. 2016). Seven of the eight regulatory variants were associated with increased growth rates relative to strains possessing the wild-type *mcr-1* regulatory sequence, providing clear evidence that the regulatory polymorphisms reduce the cost of colistin resistance (Figure.1B).

### Regulatory polymorphisms reduce *mcr-1* expression and activity

*In-silico* analysis revealed that all constructed regulatory polymorphisms are predicted to reduce *mcr-1* transcription and/or translation (Table.S2), suggesting a simple link between reduced MCR-1 abundance and increased fitness. To verify this, we took two experimental approaches to understand the link between *mcr-1* expression and fitness in the absence of colistin.

The first approach focused on MCR-1 protein activity. Mutating the catalytic site of *mcr-1* largely eliminated the cost of *mcr-1* carriage in the absence of colistin (Figure.1B), suggesting that the cost of *mcr-1* mainly comes from the activity of the MCR-1 protein. Phosphoethanolamine (pEtN) transferase activity of MCR-1 results in a reduction of membrane surface charge due to the neutralisation of negatively charged lipid A phosphate groups on lipopolysaccharide (LPS) molecules (Liu et al. 2016; Kai and Wang 2020). This reduction of cell surface charge is thought to prevent the binding of positively charged colistin to bacterial membranes (Sabnis et al. 2021). To investigate the link between MCR-1 activity and fitness, we measured the impact of regulatory mutations on cell surface charge in the absence of colistin. As expected, expressing mcr-1 with the WT regulatory sequence reduced cell surface charge, whereas this effect was partially alleviated in strains carrying regulatory variants (Figure.2A).

**Figure 2:**
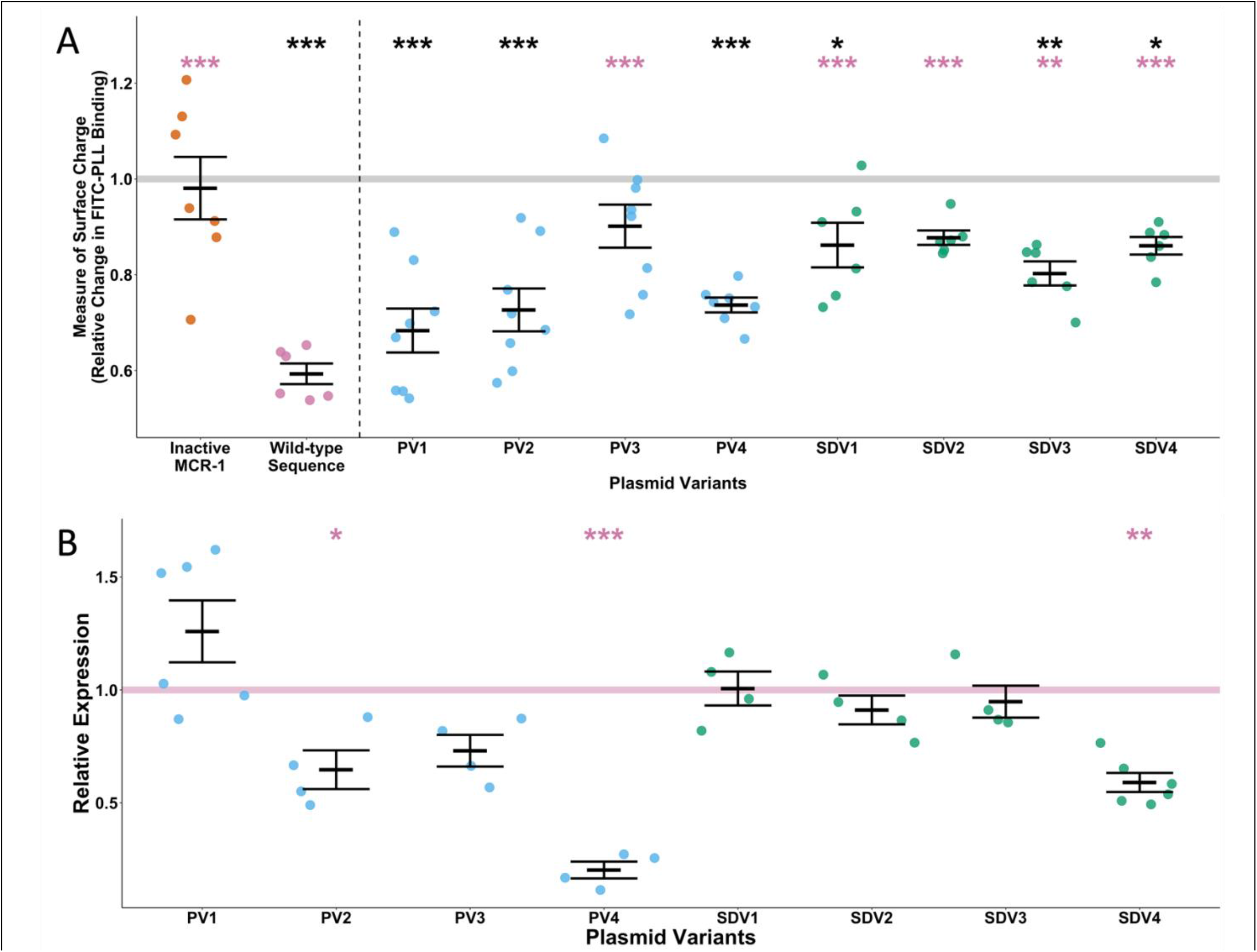
Activity and expression assessments of regulatory variants. **A**) MCR-1activity was assessed by measuring relative cell surface charge using a FITC-PLL binding assay. Surface charges of regulatory *mcr-1* variants (PV = blue, SDV= green), inactivated *mcr-1* (orange) WT sequence (pink) were measured relative to an empty control (grey line, set to 1). Significance in comparison to the empty vector (black) and consensus expression (pink) are indicated above the respective plasmid variants (p-values: ***<0.001, **<0.01, *<0.05, Dunnett’s t-test adjusted for multiple comparisons, n=6-10, error bars = SE) **B**) *mcr-1* transcript levels of regulatory variants (PV = blue, SDV= green) are shown relative the WT regulatory sequence (pink line, set to 1). Solid variant lines show mean relative expression, points individual values. Significance in comparison to consensus are indicated above the respective plasmid variants (p-values: ***<0.001, **<0.01, *<0.05, ns=not significant. Dunnett’s t-test adjusted for multiple comparisons, n = 4-6, error bars = SE)

As a second approach, we measured the impact of regulatory variants on levels of *mcr-1* transcription (Figure.2B). As expected, regulatory variants generally reduced levels of *mcr-1* transcripts which is indicative of reduced *mcr-1* transcription and/or increased mRNA degradation. This reduction in expression was particularly evident for regulatory variants with mutations in the *mcr-1* promoter region (i.e., the -10 sequence), consistent with the idea that the cause of decreased expression is reduced transcription initiation.

### Regulatory polymorphisms increase colistin resistance

Given that regulatory variants were associated with decreased *mcr-1* expression and/or MCR-1 activity, we hypothesised that the increased fitness of these variants in the absence of colistin was associated with a trade-off in terms of decreased fitness in the presence of colistin (San Millan et al. 2017; Yao et al. 2022). As a preliminary test of this idea, we measured the impact of regulatory mutations on colistin resistance using MIC assays. Surprisingly, we found that the colistin MIC of regulatory mutants were equal to or greater than that of the WT regulatory sequence (Figure.3A).

**Figure 3:**
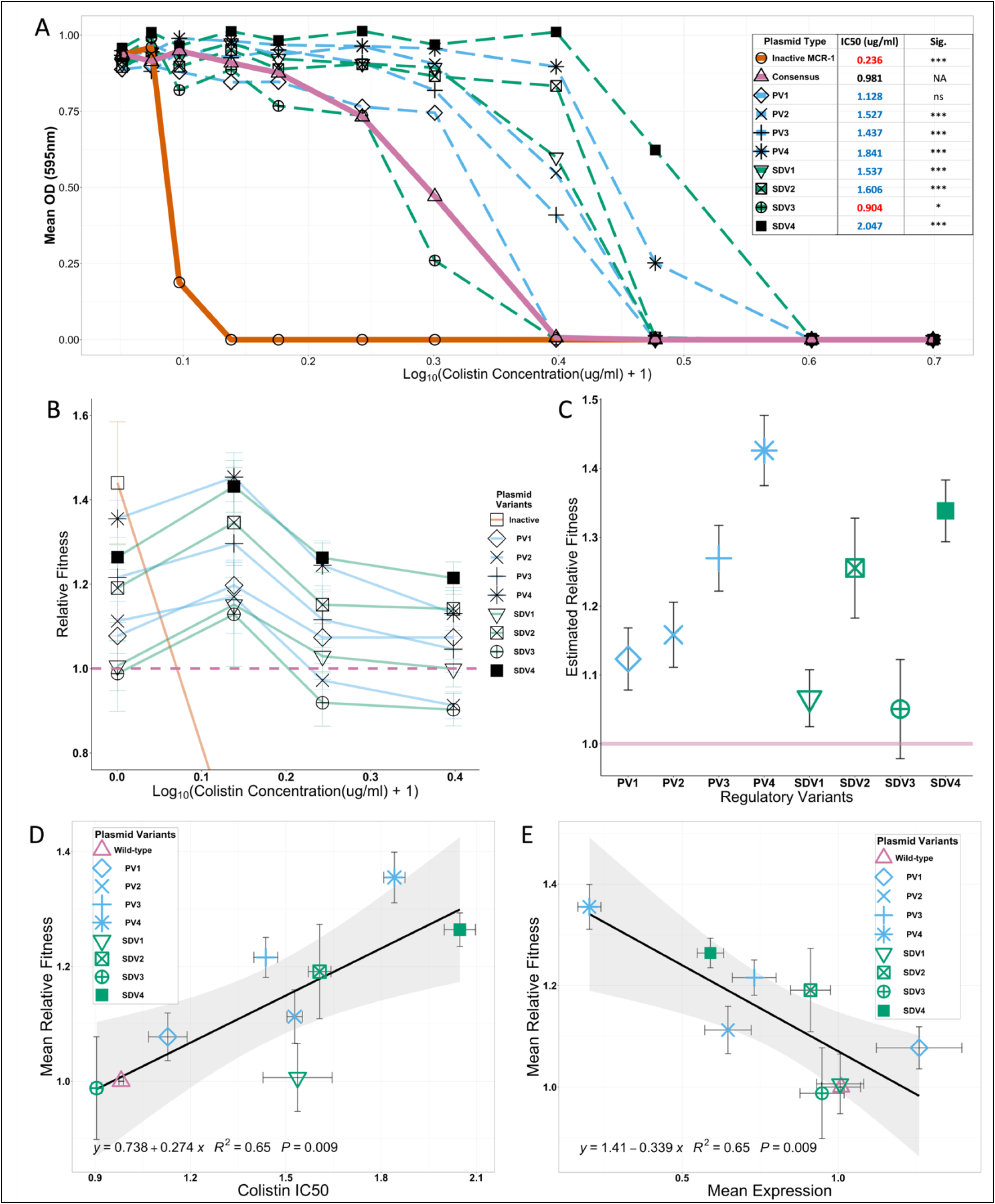
Resistance and fitness assessments of regulatory variants. **A**) Mean OD in a range of colistin concentrations (0 – 2ug/ml, Log_10_ Scale) is shown for regulatory variants (dashed lines, PV=blue, SDV=green) and controls (solid lines, pink=WT, grey=Empty control), n=15. Estimated colistin IC50s of control and variants are shown, as determined by fitting OD data to a dose-response model. IC50 values that are lower and higher than the WT highlighted in red and blue respectively, with statistical significance determined by pairwise comparisons of variants to the WT regulatory sequence (p-values adjusted using the Bonferroni correction for multiple comparisons ****<0.001, *<0.05). **B**) Relative fitness of regulatory variants and inactivated control compared to the WT control (set at 1, dotted pink line). Fitness values below the lower limit (0.8) are not shown. Error bars = standard error, n = 5. **C**) Estimated single fitness values for each regulatory variant are plotted (error bars=SE). **D-E**) Scatterplots showing the relationship between **D**) Colistin IC50 (x-axis) or **E**) mean expression (x-axis) against mean relative fitness in the absence of colistin (y-axis). Points assigned to variants/controls are indicated in the legend (error bars = SE). Linear regression model (black line) was fitted to the data (confidence intervals in grey shading). Regression line equation, model r-squared and p-value indicated in black text.

As a second approach to measure colistin resistance levels, we calculated the IC50 for colistin, which provides a quantitative estimate of the colistin concentration needed to reduce bacterial OD by 50%. Six of the regulatory variants were associated with an increased IC50, and a single variant caused a slight reduction in IC50.

As a final test of the link between *mcr-1* expression and resistance, we manipulated *mcr-1* expression by inserting the *mcr-1* gene under the control of a wild-type regulatory sequence into the *E. coli* chromosome (Figure.S1). This manipulation decreases *mcr-1* expression by moving the gene from a multi-copy pSEVA plasmid replicon (4-5 copies/cell) to a low copy number chromosomal replicon. Consistent with our earlier results, *E. coli* with a chromosomally integrated *mcr-1* had increased growth rate (Figure.S1A) and colistin resistance (Figure.S1B) compared to the positive control strains in which *mcr-1* was expressed from a low copy number plasmid.

### Regulatory fine-tuning increases colistin resistance and fitness

To further examine fitness trade-offs associated with regulatory mutations, we directly measured the impact of regulatory mutations on competitive fitness across a gradient of colistin concentrations by co-culturing *E. coli* possessing regulatory variants and WT control strains (Figure.3B).

As expected, the strain carrying catalytically inactivated MCR-1 under the control of a wild-type regulatory sequence had high fitness in the absence of colistin and low fitness in the presence of colistin (Figure.3B). Regulatory variants, on the other hand, were associated with increased fitness in both the presence and absence of colistin, although there was significant variation in fitness between regulatory variants (F-value_(7)_ = 15.3, p-value= 1.84e-14). No significant interaction was found between relative fitness of regulatory variants and colistin concentration (F-value_(21)_ = 0.44, p-value= 0.98). Given this, we were able to estimate a single fitness value for each regulatory mutant, and all regulatory mutants had higher fitness than the wild-type strain (Figure 3C). To better understand the surprising finding that *mcr-1* regulatory mutations increase strain fitness and colistin resistance, we used a linear regression to test for a correlation between resistance and fitness. In this analysis, we used only fitness data that was collected in colistin free media to avoid spurious correlations between fitness and colistin resistance. Fitness was strongly correlated with increased resistance, highlighting the ability of regulatory mutations to fine-tune *mcr-1* expression without any associated trade-offs (Figure 3D; F-value_(1,7)_=12.96, P=0.009, r^2^=.65)

Given that regulatory polymorphisms impact both *mcr-1* activity and expression, we used multiple regression to independently estimate the contribution of decreased activity and expression to increased fitness. Reduced *mcr-1* expression is correlated to fitness, demonstrating that variation in fitness between the regulatory mutants partially explains the effect that the introduced mutations have on *mcr-1* expression (Figure 3E; F-value_(1,7)_=12.8, P=0.009, r^2^=.65). The high fitness of the catalytically inactive MCR-1 mutant compared to the WT reference strain implies that MCR-1 activity carries fitness costs. However, measurements of the MCR-1 activity of the regulatory variant strains (as measured by cell surface charge) were not correlated with fitness, (Figure S2; F-value_(1,7)_=0.78, P=0.4, r^2^=.10), implying that variation in MCR-1 activity was not a consistent source of variation in fitness between the regulatory mutants.

### Evolutionary origins of regulatory variants

To better understand the evolutionary trajectory of *mcr-1* regulation, we searched for regulatory polymorphisms in a recently published genomic dataset from large-scale surveillance of colistin resistant *E. coli* from human, environmental, and agricultural sources in China from 2016-2018 (Shen *et al*., 2020, see Methods).

*mcr-1* was initially acquired by *E. coli* as a part of a composite IS*Apl1* transposon, but the IS*Apl1* elements flanking *mcr-1* degenerated following transposition to different plasmid backgrounds. This degeneration results in the loss of active transposition and a ‘fossilization’ of the *mcr-1* cassette in these backgrounds (Snesrud et al. 2016; R. Wang et al. 2018). The presence of repetitive sequences in the IS*Apl1* copies flanking *mcr-1* leads to fragmented *mcr-1* plasmid assemblies when isolates are sequenced using short-read technologies. The wild-type regulatory sequence was most associated with contigs that carried no plasmid replicon, suggesting that the WT regulatory sequence was linked to IS*Apl1* copies that fragment plasmid assemblies(Figure 4A). These contigs were shorter on average than contigs associated with regulatory variants (median 27.0 kb vs. 32.6 kb for non-wildtype sequences), providing further evidence to support this idea. Anecdotally, the presence of *mcr-1* on short contigs (<3kb) due to fragmented assembly can be evidence of the intact IS*Apl1* composite transposon. The wild-type regulatory sequence was strongly associated with such short contigs (39.4% of wild-type regulatory sequences were on such contigs vs. 12.0% for regulatory variants, n=569 isolates, Chi-squared test X^2^=55.9, p<0.001), consistent with regulatory variants being associated with ‘fossilized’ *mcr-1* sequences that lack copies of IS*Apl1*.

**Figure 4.**
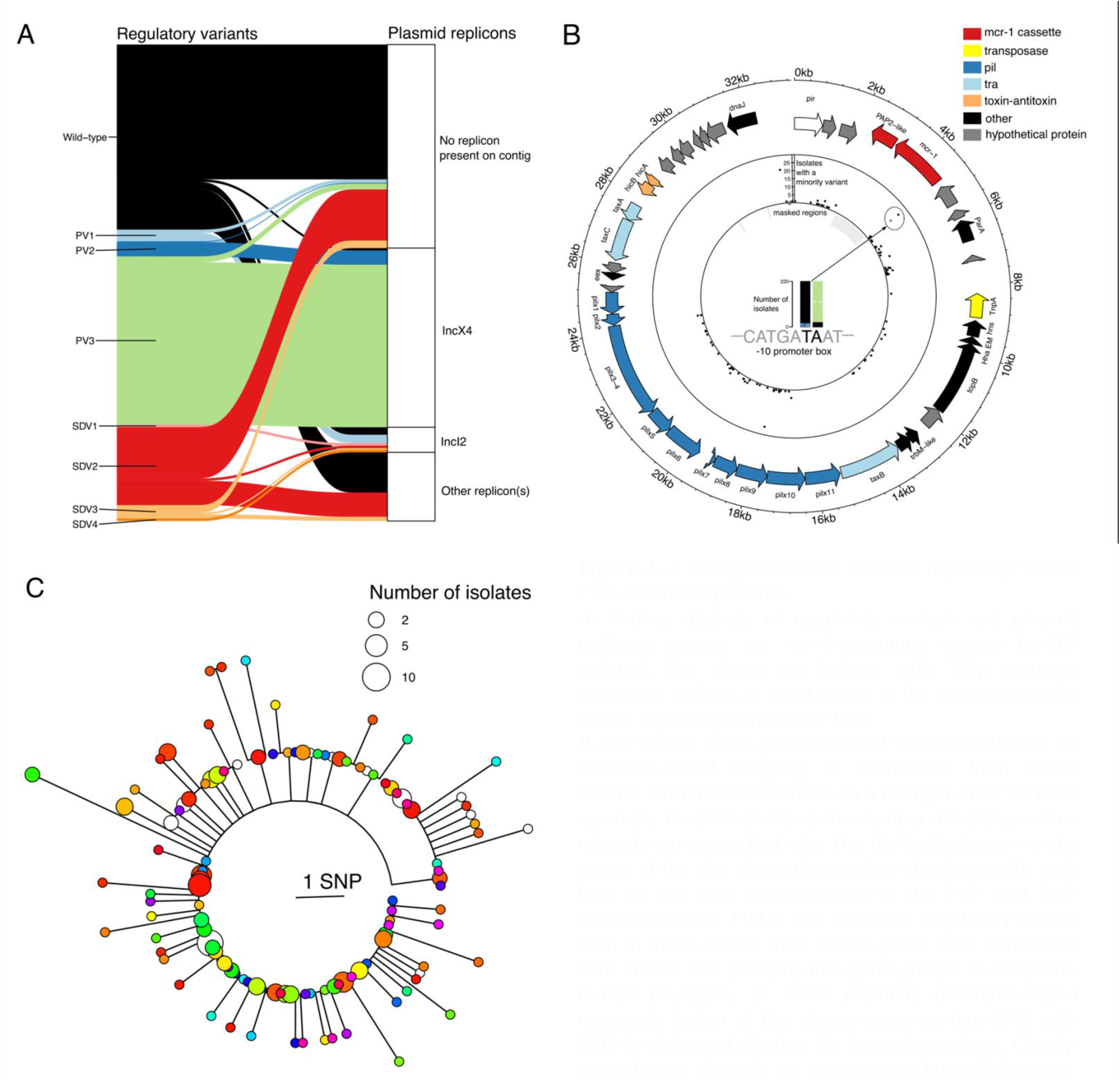
A strong association between regulatory variant PV3 and IncX4 plasmids. **A**. Sankey diagram of regulatory variants and plasmid replicons present on *mcr-1*-containing contigs (n=569 isolates). Not shown are isolates with 75bp upstream sequences without an exact match to the consensus or the named regulatory variants (n=110). **B**. The 32.6kb IncX4 plasmid (NCBI KU761327.1) used for reference-based mapping of short-reads from n=220 isolates with *mcr-1* and lncX4 on a contig in their de novo assembly. Inner track shows the number of isolates with a minority variant at that site. The highlighted zoom in the centre of the plot shows the number of isolates with each base at the sites corresponding to the PV2- and PV3-associated SNPs. PV3 is the dominant regulatory variant seen in IncX4. Outer track shows genes after annotation with Prokka and grouping into gene groups. **C**. Midpoint-rooted phylogeny of IncX4 plasmids (reference-based mapping). Colour of tips shows sequence type (ST), with ST10 in white and all other STs in random colours. Closely-related IncX4 plasmids are seen across a huge ST diversity.

*mcr-1* is almost always carried on conjugative plasmids, with the main replicons being IncX4, IncI2 and IncHI2 (Sun et al. 2018; C. Shen et al. 2020). We observed strong associations between regulatory variants and plasmid replicons (Figure 4A). Most notably, there was a strong association between IncX4 and regulatory variant PV3: 194/220 (88.2%) of IncX4-containing contigs also had PV3, and 194/201 (96.5%) PV3-containing contigs also had IncX4. No IncX4-containing *mcr-1*-containing contigs had an intact copy of IS*Apl1*, supporting the idea that the *mcr-1* cassette is no longer transposable in IncX4 backgrounds and the spread of *mcr-1* with the PV3 mutation is mediated by transfer of IncX4 plasmids, and not by transposition.

To better understand the process of the evolution of regulatory variants, we reconstructed a phylogeny of closely related IncX4 plasmids from human, agricultural and environmental sources using a reference IncX4 *mcr-1*-positive plasmid first sequenced in 2014 (see Methods). Although the genetic diversity of IncX4 plasmids is low, the phylogeny suggested that regulatory fine-tuning in IncX4 has occurred twice: first with PV3, and then subsequently with PV2 in a small subclade (Figure S3). The presence of compensatory mutations on a conjugative plasmid generates a worst-case scenario for resistance management, as it creates the potential for low-cost AMR plasmids to transfer between host strains (Zwanzig et al. 2019). To test the role of this mobility in the dynamics of *mcr-1*, we mapped host strains onto the IncX4 phylogeny. Plasmids with the PV3 regulatory variant were seen across a broad diversity of host strains (n=71 STs). This highlights how a conjugative plasmid with a fine-tuned resistance gene can rapidly disseminate low-cost resistance across diverse strains. While it is also possible that PV3 was independently acquired by closely related plasmids in different STs, we believe this is unlikely. The low diversity of IncX4 plasmids and the strong association between plasmid replicons and compensatory mutations suggest that repeated evolution is an unlikely model for the dissemination of the PV3 regulatory variant. The most common host strain for the IncX4 plasmid was ST10 (n=26 isolates), a known and prevalent livestock-associated extra-intestinal pathogenic *E. coli* lineage (Manges et al. 2019). This could partly explain why ST10 may be an important reservoir of plasmids that can be transferred to other strains, as suggested previously for MCR-1 (Matamoros et al. 2017).

### Regulatory mutations stabilize colistin resistance

Fitness costs are thought to play a key role in the long-term stability of antibiotic resistance, particularly if antibiotic consumption is low. To test the impact of regulatory mutations on *mcr-1* stability, we calculated changes in the prevalence of regulatory polymorphisms before (2016) and after (2017, 2018) colistin was banned as growth promoter using data from large-scale genomic surveillance projects. Regulatory variants had already achieved a high frequency by the time that colistin was banned as a growth promoter, and the frequency of regulatory variants increased over time (Figure 5A). The continued rise of regulatory variants suggest that regulatory mutations were selectively advantageous under low levels of colistin consumption. However, interpreting this increase is somewhat difficult given differences in the number of MCR positive isolates recovered from different one-health sources in different years. To overcome this difficulty, we focused on the prevalence of *mcr-1* in pigs, which were intensively sampled due to the importance of farms as a source of colistin resistance (66 farms, 684-1575 pigs per year). This revealed that the proportions of tested regulatory variants increased between 2016 and 2018 relative to WT sequences (2016 to 2018: 34/68 to 32/48, p=0.037; one-sided two-sample test for equality of proportions without continuity correction) and as a proportion of all sequenced *mcr-1*-positive isolates (34/77 to 32/58, p=0.11): one-sided two-sample test for equality of proportions without continuity correction). Crucially, carriage rates of *mcr-1* with wild-type regulatory sequences declined at approximately 1.5 times of the rate of decline of those carrying tested regulatory variants (Figure.5B).

**Figure 5:**
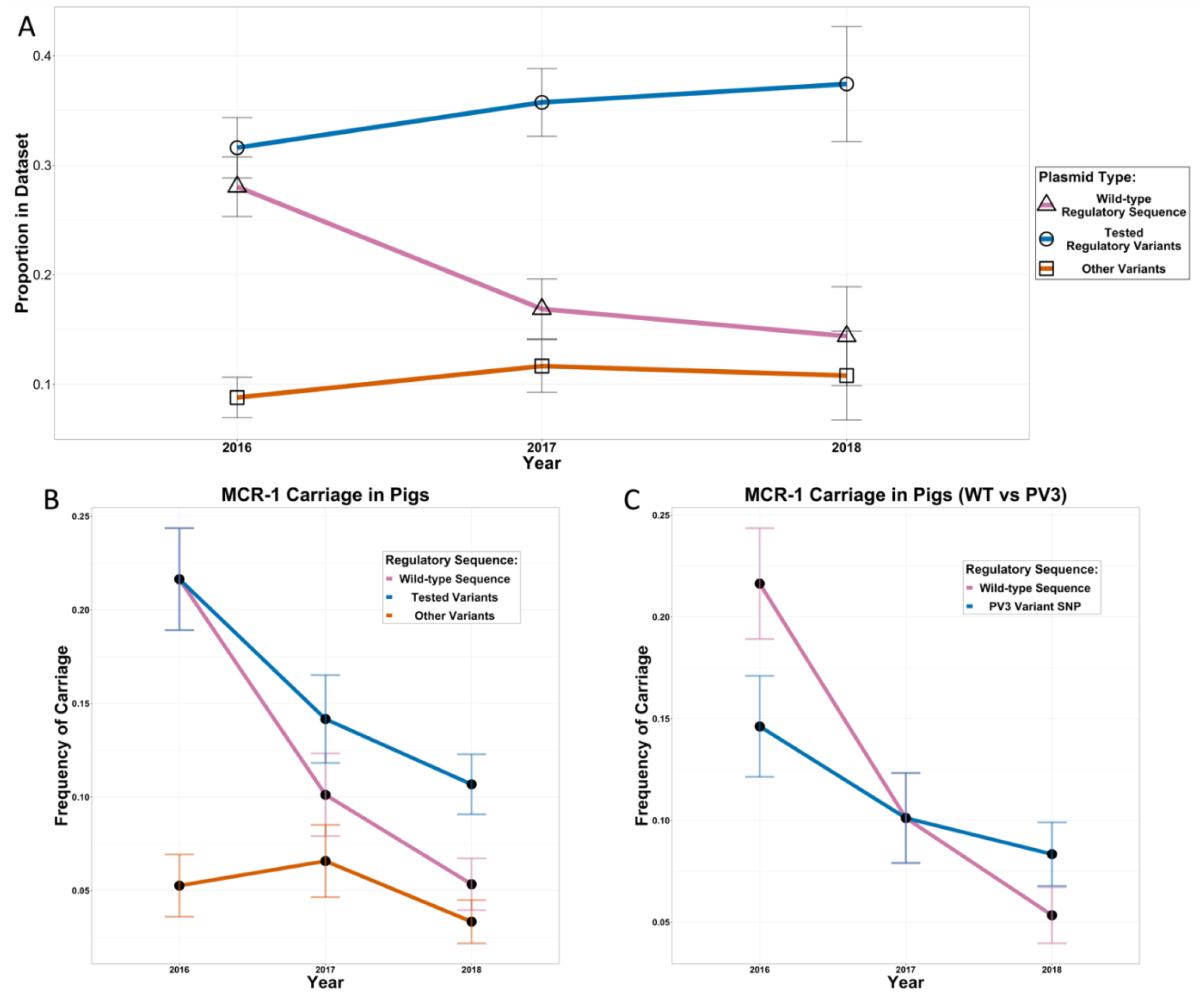
Variant prevalence and frequency in Shen sequence 2020 dataset. **A**) Proportion of wild-type (pink) and variant (blue) regulatory sequences in *mcr-1*+ sequences separated by year. Sequences carrying untested variants (orange) are also shown. Error bars show standard error of proportions. B) Frequency of carriage (i.e., Frequency of *mcr-1* prevalence * frequency of variant prevalence) was determined for wild-type (pink, slope=-0.081) and variant (blue, slope=-0.055) and untested variant (orange, slope=-0.01) sequences across the years for pig samples. Each sample was treated as an independent unit. Error bars show the propagated standard error. **C**) Frequency of carriage of wild-type (pink, slope=-0.081) sequences compared to those carrying PV3 (blue, slope=-0.031) variant SNPs. Error bars show the propagated standard error.

A closer look at the sequences revealed that most *mcr-1*+ isolates obtained post colistin ban carried IncX4 plasmids with PV3 regulatory mutations (Figure.S4). Additionally, sequences carrying the PV3 variant showed additional reductions in their rate of decline (Figure.5C). This is of note as PV3 is by far the most prevalent regulatory variant and has surpassed wild-type regulatory sequences in prevalence among *mcr-1*+ sequences post-ban (Figure.S5). After the ban on colistin, the proportion of sequenced *mcr-1*-positive isolates with PV3 increased compared to WT (2016 to 2018: 25/59 to 25/41, p=0.033; one-sided two-sample test for equality of proportions without continuity correction) and as a proportion of all sequenced *mcr-1*-positive isolates (25/77 to 25/58, p=0.102).

## Conclusion

The rapid spread of *mcr-1* mediated colistin resistance in *E. coli* represents an important threat to human health given that colistin provides a ‘last-line of defence’ for the treatment of infections caused by multi-drug resistant *E. coli*. Here we show that the *mcr-1* regulatory region has evolved to reduce the cost of *mcr-1* while simultaneously increasing colistin resistance, providing a poignant demonstration of the ability of mutation and natural selection to fine-tune gene expression. Conjugative plasmids carrying *mcr-1* promoter variants have transferred between host strains, disseminating low cost/high colistin resistance *mcr-1* across *E. coli*. Crucially, regulatory variants were associated with increased *mcr-1* stability in pigs following a ban on the use of colistin as a growth promoter that reduced colistin use in agriculture by 90%. These findings provide a clear and unambiguous example of how the adaptive evolution of resistance genes together with horizontal gene transfer can stabilize antibiotic resistance and limit the efficiency of interventions aimed at reducing antibiotic consumption.

*mcr-1* was probably mobilised from the chromosome of pig-associated bacteria by a composite transposon that subsequently transferred *mcr-1* to plasmids associated with *Enterobacteriaceae* (R. Wang et al. 2018). We speculate that the increase in gene copy number associated with moving from a chromosome to multi-copy plasmids caused *mcr-1* to be overexpressed, resulting in low levels of colistin resistance and high fitness costs. Promoter mutations that reduce *mcr-1* expression provide a simple solution to offset the impact of increased gene copy number while allowing *mcr-1* to continue to benefit from the high mobility provided by conjugative plasmids. For example, plasmid transfer between strains of *E*.*coli* with different niches may help to stabilize low cost/high resistance *mcr-1* alleles in the face of perturbations, such as altered antibiotic use, that favour a sub-set of *E*.*coli* strains (T. Wang et al. 2022). An important challenge for future work will be to directly test the importance of horizontal gene transfer in stabilizing antibiotic resistance in pathogen populations (Lopatkin et al. 2017; Bergstrom, Lipsitch, and Levin 2000). This may be especially important in a one-health context where HGT can allow resistance genes to spread across strains associated with human, agricultural and environmental niches (Hernando-Amado et al. 2019; Y. Wang et al. 2017; Y. Shen, Wu, et al. 2018).

One of the simplest interventions to combat AMR is to reduce antibiotic consumption, and the ban on the use of colistin as a growth promoter represents one of the largest and best studied attempts to combat resistance using this approach. In many respects, the evolution of *mcr-1* towards increased resistance and fitness and on-going transmission of *mcr-1* across *E. coli* strains represents a worst-case scenario for resistance management. However, it is important to emphasize that the prevalence of *mcr-1* continued to decline across one-health sectors following the ban on use of colistin as a growth promoter (C. Shen et al. 2020; Y. Wang et al. 2020). The key implication of these findings for resistance management is that the evolution and transmission of resistance genes may need to be considered to accurately forecast the impact of reducing antibiotic consumption on AMR. Our findings suggest that in extreme cases where resistance has negligible costs and/or a very high rate of transfer, limiting antibiotic consumption may not be a viable strategy to reduce resistance.

## Methods

### Strains, plasmids, and growth conditions

All experiments were carried out in E. coli MG1655 using Luria-Bertani (LB) or Mueller Hinton Broth (MHB) medium (Sigma-Aldrich). All control and constructed plasmids are listed in Supplementary table.S3. Media was supplemented with colistin Cayman Chemical) or ampicillin (Sigma-Aldrich) as appropriate to enable selected growth of plasmid carriers and for MIC assays.

### Oligonucleotides

A full list of DNA oligonucleotides used in this work is provided in Supplementary table.S4. All oligonucleotides were ordered from ThermoScientific.

### *mcr-1* Regulatory Variant Construction

A synthetic pSEVA vector containing *mcr-1* and its natural promoter as a cargo was constructed. *mcr-1* and its surrounding regions (75bp upstream, 40bp downstream) were PCR-amplified from the PN16 (IncI2) plasmid using Q5® High-Fidelity DNA Polymerase (New England BioLabs). The amplified fragment was cloned into pSEVA121 using the NEBuilder® HiFi DNA Assembly kit (New England BioLabs) as per manufacturer’s instructions. Plasmids containing one of the eight regulatory variants and inactivated (T285A) *mcr-1* were generated using mutagenic primers and the Q5® Site-Directed Mutagenesis Kit New England BioLabs) as per manufacturer’s instructions. Similar mutagenic primers were used to introduce unique qPCR sequence tags for each variant and control plasmid. Sequence verified plasmids were transformed into MG1655 *E. coli* strains.

### Construction of chromosomally Integrated *mcr-1*

MG1655 with chromosomally integrated *mcr-1* was constructed by lambda red recombineering. The consensus *mcr-1* gene and regulatory sequences were cloned into the MG1655 chromosome in replacement of the non-essential lacZ gene. Isolates were confirmed for correct assemblies using blue-white screening and sequence verification.

### Resistance determination via minimum inhibitory concentration (MIC) assays

Colistin resistance of constructed strains was determined using standard broth microdilution methods and OD measurements. Bacteria were grown in MHB media supplemented with the appropriate antibiotics (50 µg/ml ampicillin for MG1655 containing pSEVA plasmids and 1 µg/ml colistin for MG1655 containing chromosomally integrated *mcr-1*). Bacteria were diluted to 5×105 CFU/ml in a range of colistin concentrations (Cayman Chemical, 0-8ug/ml) in alternating two-fold dilutions. Eight independent replicates were performed for all strains and concentrations. Bacteria were grown in 96-well culture plates overnight (37°C, 250 rpm). OD measurements (595nm) were taken following incubation with subtraction of media background measurements. OD values below 0.1 were determined to have no bacterial growth with the lowest concentration where this was reached determined as the MIC. IC50 values were estimated by fitting dose response curve (DRC) models to MIC data using the “drm” package (v2.5-12).

### Growth Rate Measurements

Overnight cultures of constructed and/or control MG1655 were diluted to 5×105 CFU/ml in MH media. Bacteria were incubated overnight at 37°C with OD measurements (595nm) taken every 10 minutes. Regression lines were fitted onto the exponential phases of growth curves. Exponential growth rate (mOD/min) was determined as the maximum gradient obtained over 10 OD measurement points. Experiments were repeated 5 times per variant on 8 different days. Growth rates varied systematically between assays carried out on different days, as judged by an ANOVA including a main effect of assay day, and we corrected for this by using residual growth rates after correcting for the effect of assay day.

### Gene Expression Easements using RT-qPCR

*mcr-1* expression levels were quantified in MG1655 containing control or regulatory variant plasmids. Total RNA was extracted from bacteria following incubation in MH supplemented with Ampicillin (50ug/ml). Bacteria were sub-cultured, grown to an OD of 0.5 (595nm) and washed in PBS. Bacterial digestion was performed using RNAprotect Bacteria Reagent (Qiagen) as per manufacturer’s instructions. Digested bacteria were subjected to RNA extraction using the RNeasy Mini Kit (Qiagen) and QiaCube liquid handling platform (Qiagen) as per manufacturer’s instructions. Extracted RNA was DNAse treated using the TURBO DNA-free kit (ThermoFisher) and quantified using the Quantifluor® RNA system (Promega).

Extracted RNA was diluted to 5ug/ml and subjected to one step RT-qPCR on the StepOnePlus(tm) Real-Time PCR System (Applied Biosystems) using the Luna® Universal One-Step RT-qPCR Kit (New England Biolabs). *mcr-1* and trfA (reference) expression were quantified using appropriate primers (Supplementary table 4). trfA was selected as a reference due to its pSEVA121 localisation allowing normalisation of plasmid copy number across all control and regulatory variant plasmids tested. Relative *mcr-1* expression was calculated using the delta-delta (2–ΔΔCt) Ct method where expression levels of variant regulatory sequences were compared to the consensus.

### Bacterial Surface Charge Measurements

Fluorescein isothiocyanate-labelled poly-L-lysine (FITC-PLL, Sigma) binding assays were used to determine *mcr-1* activity. Positively charged FITC-PLL can bind gram-negative outer membrane in a charge dependant manner due to negative charges on lipid A (Rossetti et al., 2004). *mcr-1* activity reduces these negative charges allowing estimations of *mcr-1* activity by measuring cell fluorescence. Overnight cultures of constructed and/or control MG1655 were washed and 1X Phosphate-buffered saline (PBS) buffer to a final OD595 of 0.1. FITC-PLL solution was added to re-suspended cells (5 µg/ml) and samples were incubated at room temperature for 12 min. Following centrifugation (6000xg, 5 minutes), fluorescence measurements (Ex-500nm/Em-530nm) of supernatants were compared with PBS controls to determine the proportion of bacterial-bound dye.

### qPCR Fitness Competitions

Primers were designed to match unique sequence tags on control and variant plasmids to determine plasmid concentrations via specific amplification (Supplementary Table xx). Specificity and optimisation tests were performed for all primers using PCR and qPCR. For competitions, overnight MG1655 cultures carrying control or regulatory variant pSEVA plasmids were OD normalized and diluted to pooled mixes (5×10^5^ CFU/ml total). Mixes were used to inoculate MH media of 4 conditions (0, 0.375, 0.75 and 1.5ug/ml colistin) and competitions were carried out over a 20hr incubation at 37°C. Four independent replicates were performed per competition. Resultant competition media and inoculation mixes were boiled (100^°^C, 20 minutes) and qPCR was carried out using the StepOnePlus Real-Time PCR System (Applied Biosystems). Variant specific primers were used to obtain plasmid specific CTs. Standard curves were generated for each primer pair to allow determination of variant plasmid concentrations in each competition.

Relative fitness was calculated for each condition using the following equation.

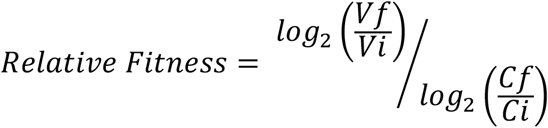

Where Vf = variant plasmid concentration following competition, Vi = variant plasmid concentration of initial pools, Cf = consensus *mcr-1* plasmid concentration following competition and Ci = consensus *mcr-1* plasmid concentration of initial pools. An ANOVA was fitted to the fitness data including effects of variant, colistin concentration (categorical variable or continuous variable) and a variant*colistin concentration interaction term. Note that this model did not include data from the variant with a catalytically inactivated MCR-1.

Linear regression models were fitted to variant relative fitness vs colistin concentration data. Y-intercepts of the resulting models were used as single estimated fitness values for the different regulatory variants.

### Genomic datasets

#### Dataset

We downloaded n=688 paired-end short-read sequencing datasets for *mcr-1*-positive *E. coli* isolates from Shen et al. 2020 (NCBI BioProject PRJNA593695). These isolates were collected between 2016-2018 from pigs, humans (both healthy volunteers and hospital inpatients), food and the environment in Guangzhou, China. We trimmed adapters with Trimmomatic v0.39 then de novo assembled isolates with Spades v3.15.3 (-k 21,33,55,77, otherwise default parameters). Processed data and scripts are available on figshare.

#### Analysing regulatory variants

We wrote custom python scripts to extract *mcr-1*-containing regions from de novo assemblies, detect plasmid replicons using ABRicate, and classify variants in the promoter region. These scripts are packaged as the conda package ‘mcroni’ (see: https://github.com/liampshaw/mcroni). To assign regulatory variants, we required an exact match to the 75bp upstream sequences used in experiments (consensus and the eight named variants: PV1-4 and SDV1-4.). Sequences that did not exactly match a named sequence were categorised as “other” (n=110 from de novo assemblies).

#### IncX4 plasmid analysis

After de novo assembly and analysis with mcroni, we selected n=220 isolates which had an IncX4 replicon on their *mcr-1*-containing contig. The median length of these contigs was 32,643 bp (IQR: 32,641-32,850 bp, range: 9,331-40,975 bp). We then used a reference-mapping approach to construct a phylogeny of closely related IncX4 plasmids. To select a suitable reference plasmid, we searched PLSDB v2021_06_23_v2 using a de novo assembled 32.6kb plasmid from isolate SRR15732044. We chose an *mcr-1*-containing IncX4 plasmid (mash similarity >0.998) isolated from a *K. pneumoniae* isolate from peritoneal fluid from a hospital inpatient in China in September 2015 (KU761327.1). The same study also found an identical plasmid in an *E. coli* isolate from a separate patient in August 2014. We mapped reads to this reference with snippy v4.6.0 (--mincov 10, --minfrac 0.9, otherwise default parameters) which uses bwa and Freebayes. We then used snippy-core to identify core SNPs, masking a region containing the remnants of the *mcr-1*-containing cassette from the start of *mcr-1* i.e., not including the upstream *mcr-1* regulatory region (bases 2337-4762 inclusive; positions with respect to KU761327.1) and two 152bp regions with a repeated CDS which meant that reads did not map uniquely (29959-30111, 32039-32191). This repeated CDS had a blast hit to a YajB protein (AIF97194.1; e-score=4e-14, 88% query cover, 77.27% identity).

Nine isolates had unaligned positions (range: 9-10,512 unaligned sites after masking) which could indicate either incomplete sequencing or different IncX4 backgrounds. We removed these isolates leaving n=211 isolates. Consensus plasmid sequences from mapped reads had a median of 1 SNP against the reference IncX4 plasmid (range: 0-4). This corresponds to an estimated mutation accumulation rate of 0.3 substitutions per plasmid per year, or ∼1e-5 substitutions per site per year. For plotting, we used ggplot2 v3.3.6 and cowplot v1.1.1, ggsankey v0.0.99999 (Figure 4A), circlize v0.4.15 (Figure 4B), and ggtree v3.2.1 (Figure 4C, Supplementary Figure S3). Data and scripts for the re-analysis of the Shen et al. (2020) datasets are available on figshare (doi: 10.6084/m9.figshare.20943256).

#### Frequency of Carriage Calculations

Frequency of carriage of variant and consensus sequences were obtained by multiplying frequency of *mcr-1* prevalence with frequency of variant prevalence. *mcr-1* prevalence data in pigs and hospitalised patients were obtained from tables 1 and 2 respectively. Sequence data from Shen 2022 was analysed to obtain the frequency of variant prevalence. 95% confidence intervals were obtained for both frequencies. The propagated uncertainty for multiplication was calculated using the following equation:

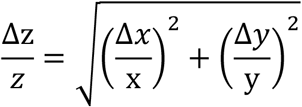

Standard error values were obtained from 95% confidence intervals using the following equation:

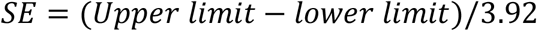

Data was visualised using ggplot. We tested for a difference in the frequency of variants by comparing the proportion of isolates with a designated variant to the proportion of isolates with a wild-type regulatory sequence using the normal approximation to the binomial. We compared regulatory variant frequency for pig populations in 2016 (pre-ban) to the frequency in 2018 (post-ban) as a proportion of all tested regulatory sequences and as a proportion of all sequenced mcr-1 positive pig isolates. We also compared the PV3 variant frequency for pig populations in 2016 (pre-ban) to the frequency in 2018 (post-ban) as a proportion of PV3 and WT sequences and as a proportion of all sequenced mcr-1 positive pig isolates. Two-sample tests for equality of proportions were conducted one-sided and without Yate’s correction for continuity.

## Acknowledgements and Funding

This work was supported by grants from the Wellcome Trust (106918/Z/15Z, C.M.) and the Medical Research Council (MR/S013768/1, T.W. and C.M.). L.O. was supported by the Biotechnology and Biological Sciences Research Council doctoral training partnership (BB/M011224/1). The computational aspects of this research were supported by the Wellcome Trust Core Award Grant Number 203141/Z/16/Z and the NIHR Oxford BRC. The views expressed are those of the author(s) and not necessarily those of the NHS, the NIHR or the Department of Health. LPS is a Sir Henry Wellcome Postdoctoral Fellow funded by Wellcome (grant 220422/Z/20/Z). We are grateful to Guo-Bao Tian and Cong Shen for helpful correspondence regarding data from their study and for making this data available via NCBI.

## Supplementary Figures/Tables

**Supplementary table 1:**
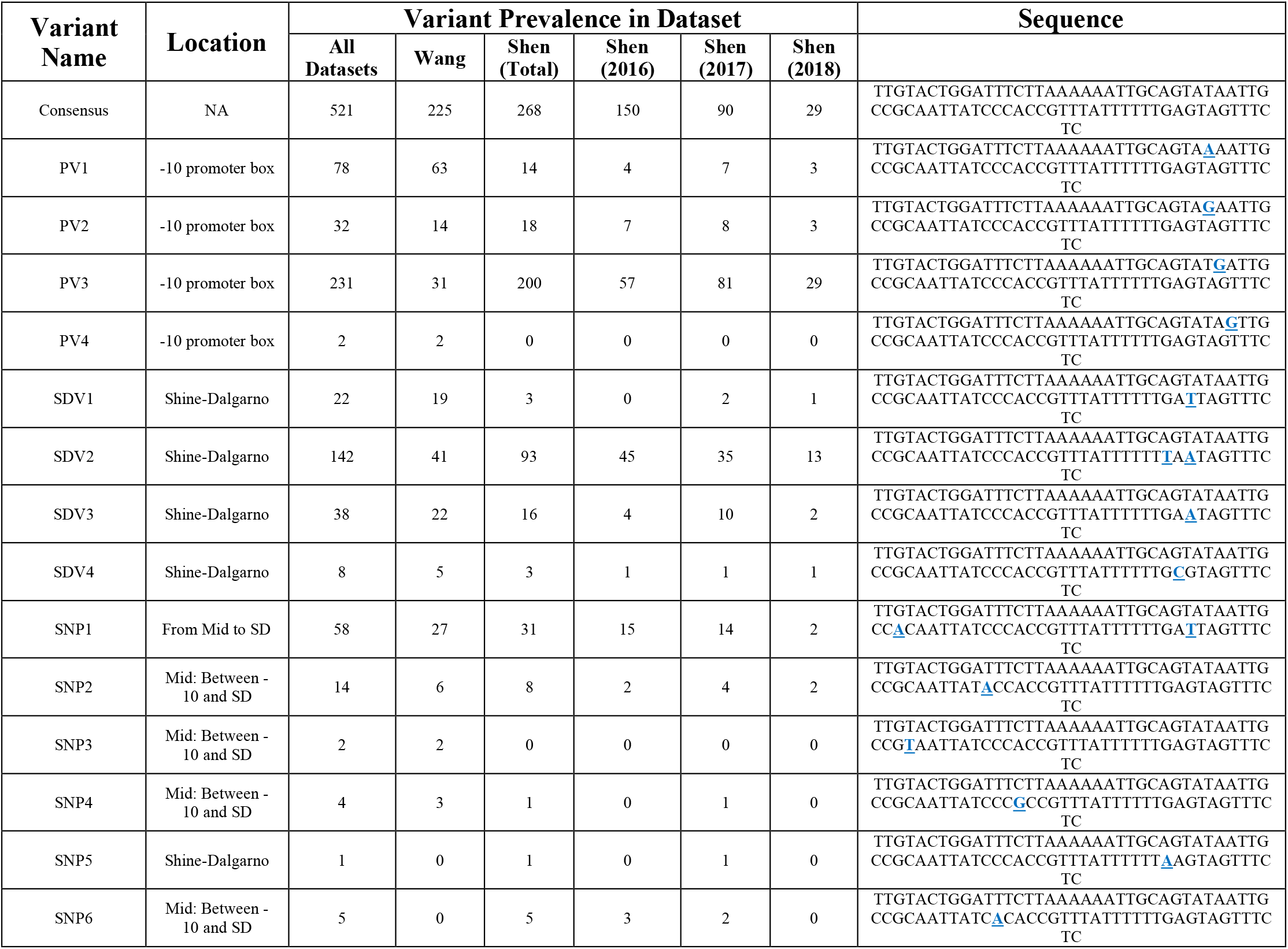
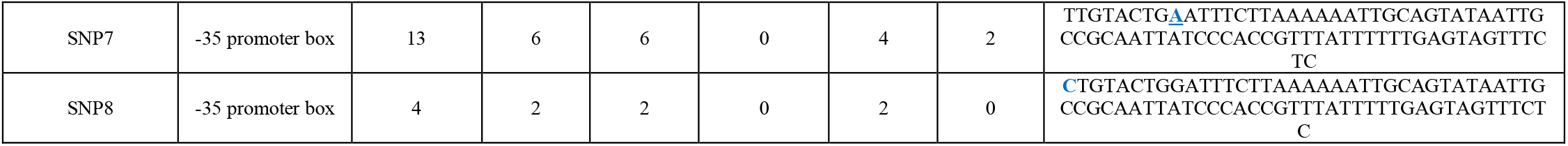
Counts of consensus and variant sequences in Shen and Wang datasets. Variant sequences are listed with variants highlighted (blue) in bold underline.

**Supplementary table 2:**
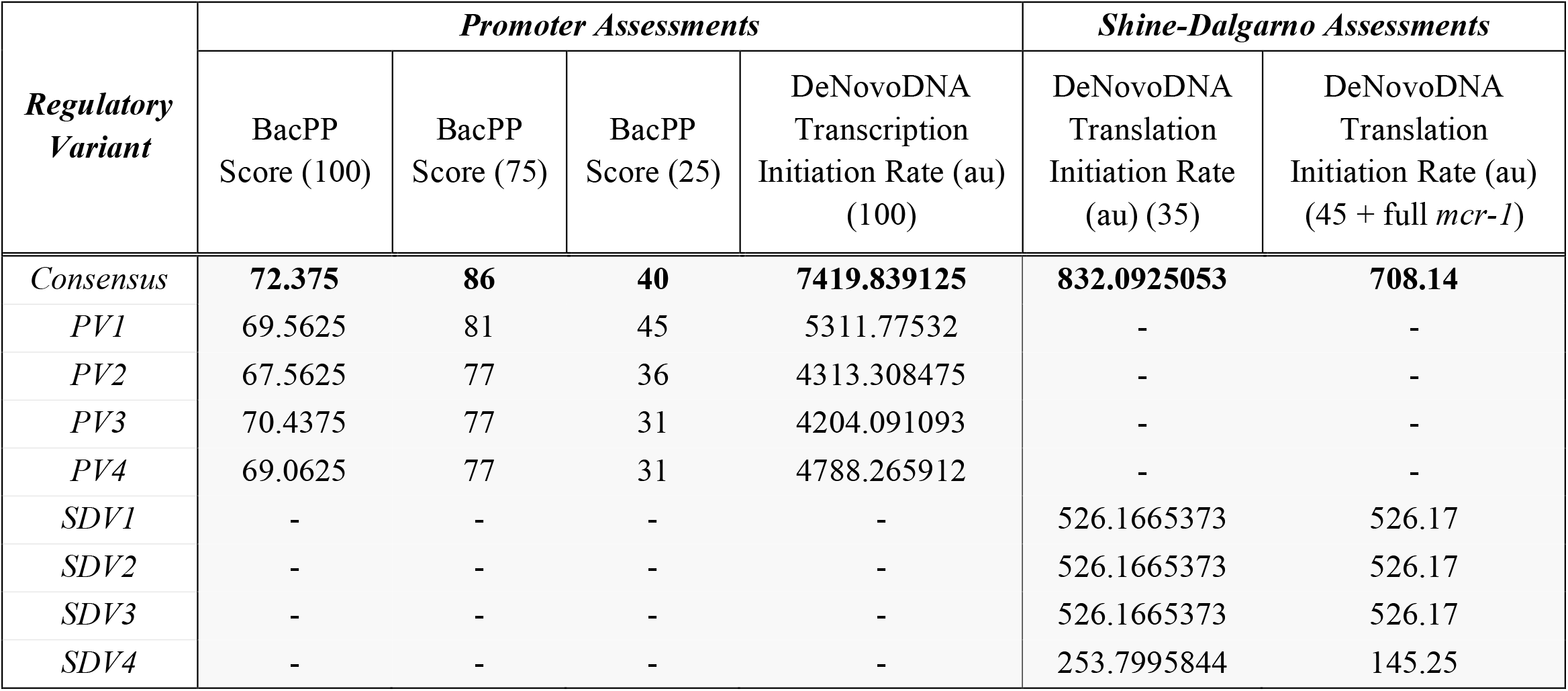
Scores obtained for promoter and Shine-Dalgarno using In-Silico analysis. Numbers in brackets indicate query length (in base pairs).

**Supplementary table 3:**
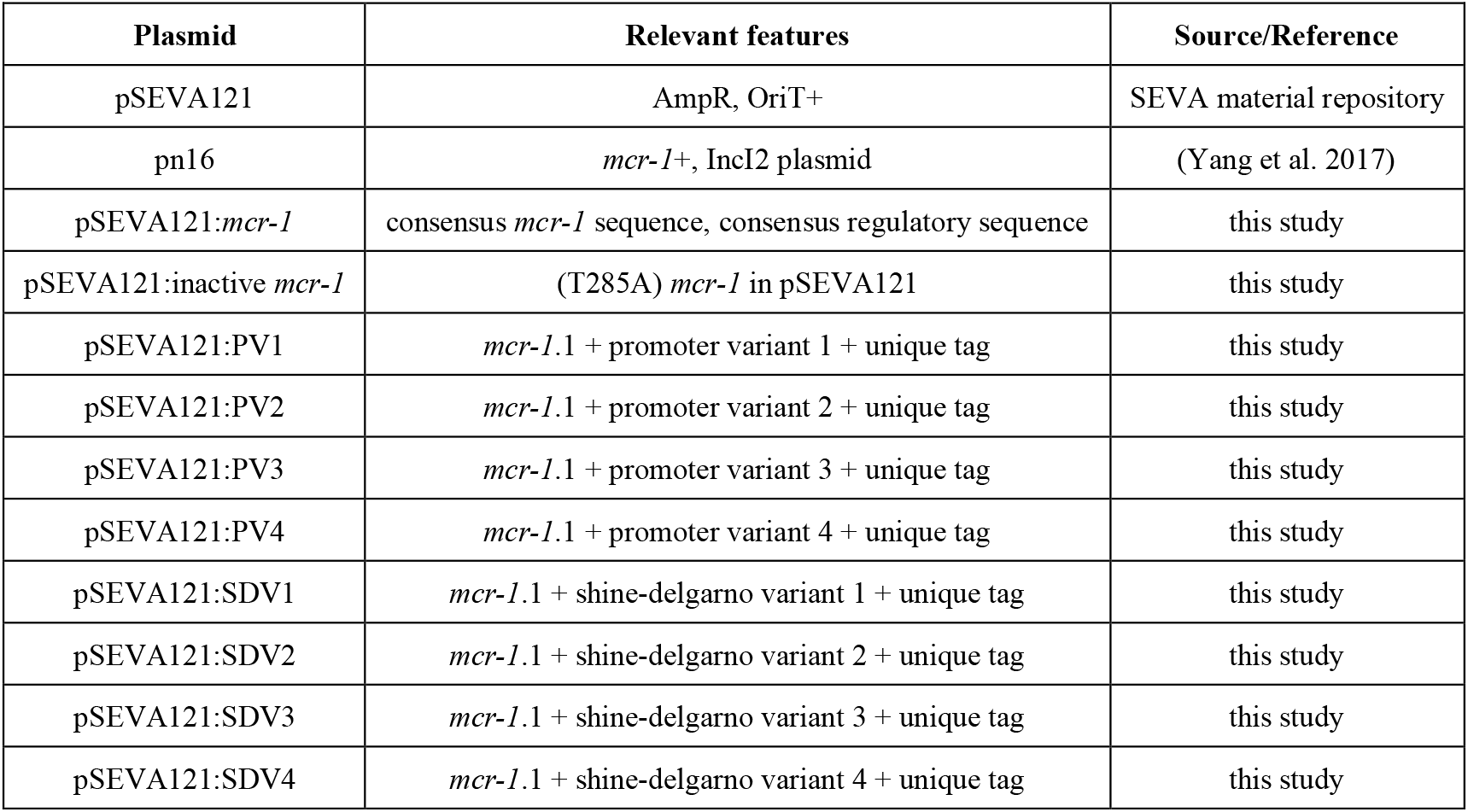
List of plasmids used/created in this study

**Supplementary table 4:**
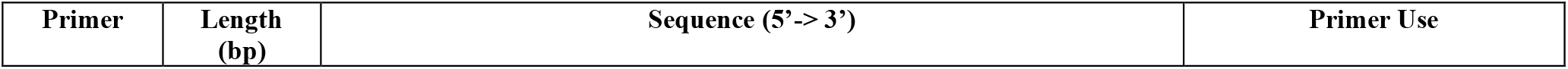
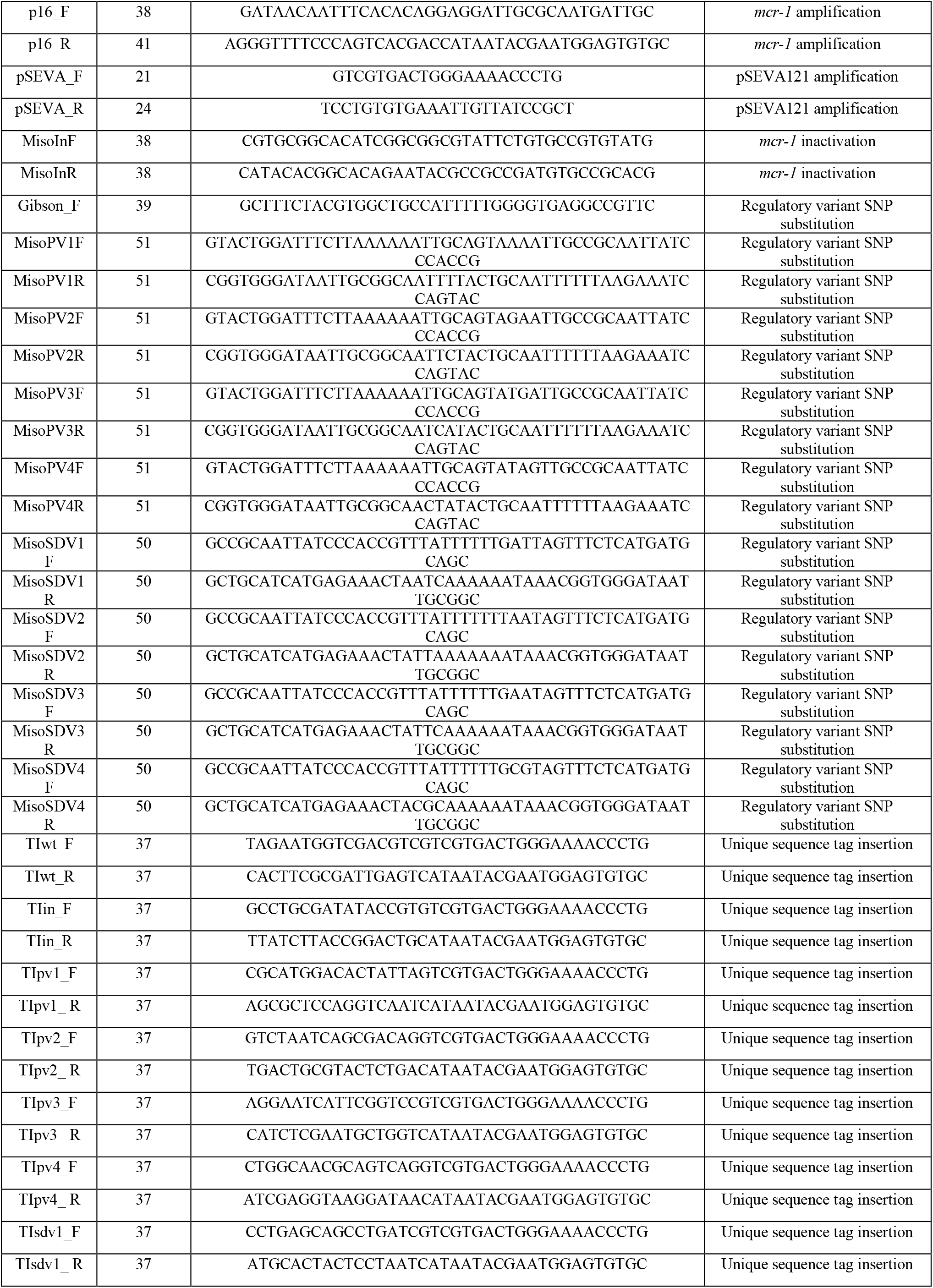
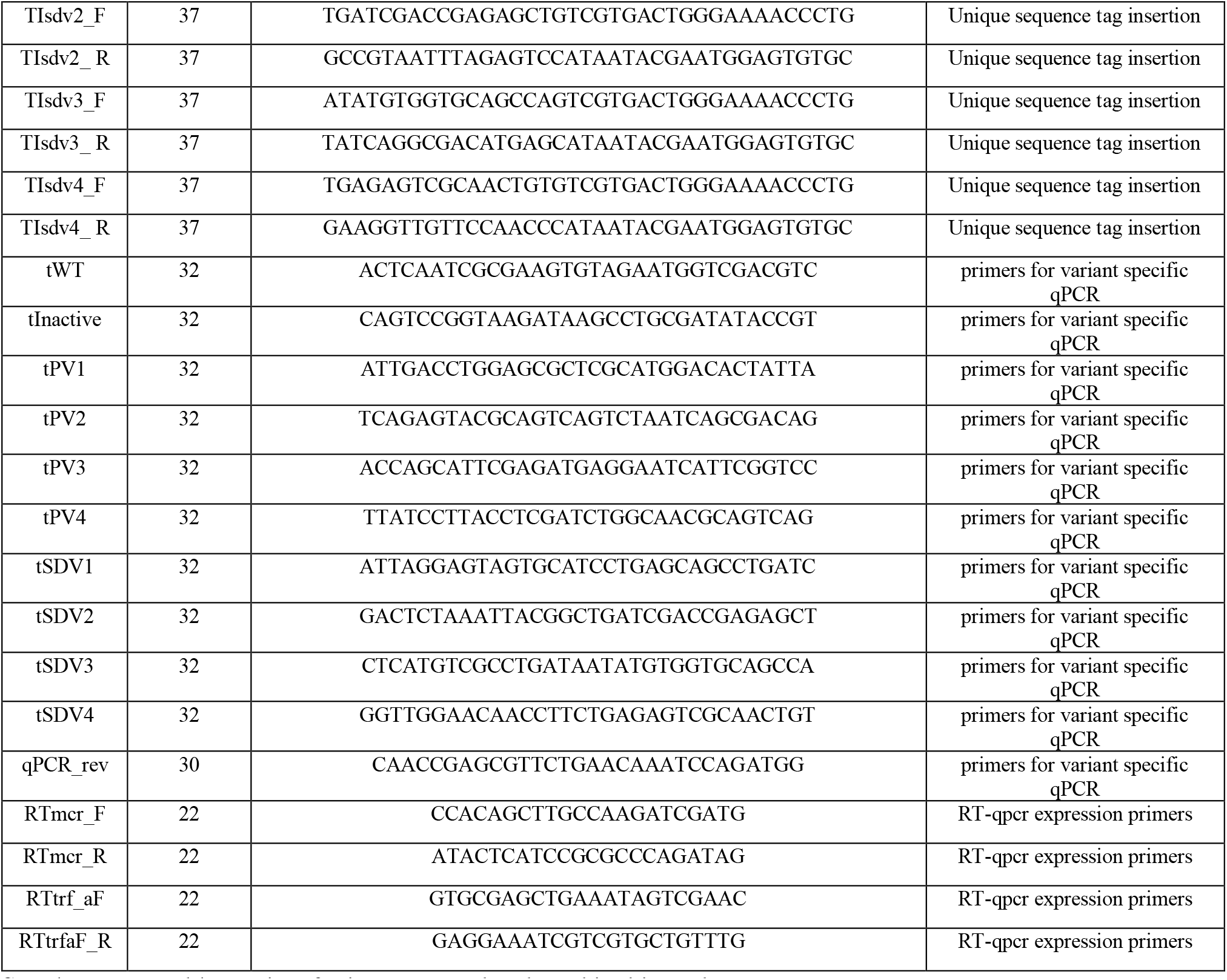
List of primers created and used in this study

**Supplementary Figure S1:**
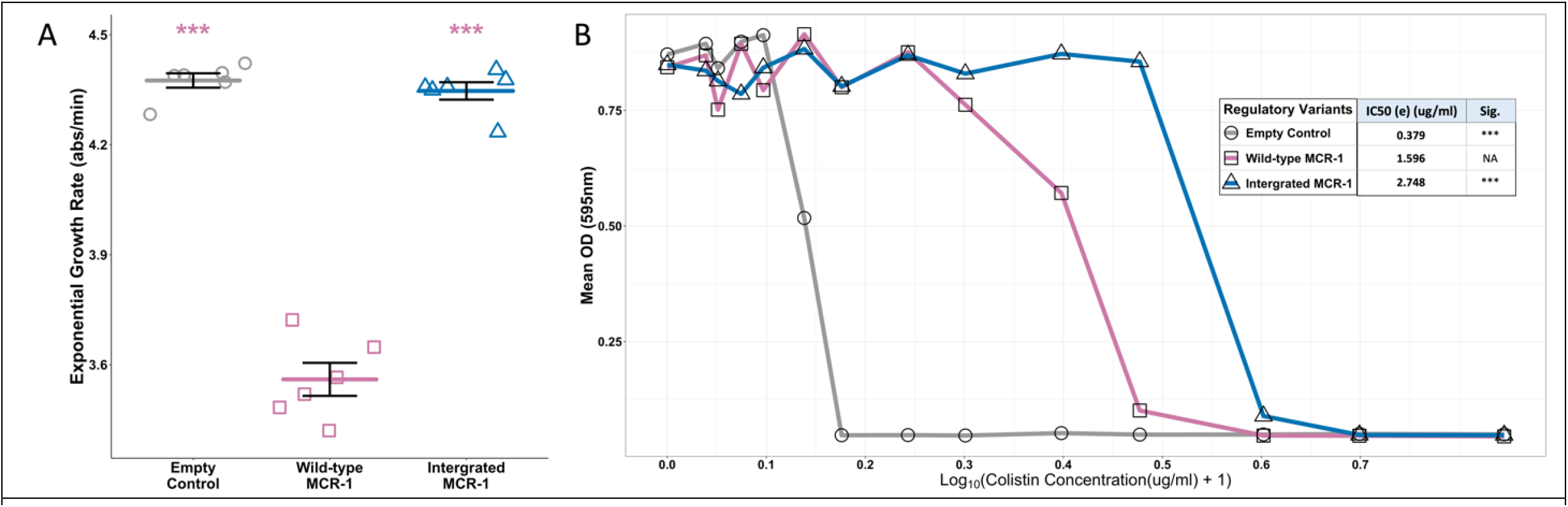
Assessments of Integrated *mcr-1* E. coli. **A)** Mean OD in a range of colistin concentrations (0 – 6ug/ml, Log_10_ Scale) is indicated for empty vector (grey) plasmid *mcr-1* (pink) and chromosomally integrated *mcr-1* (dark blue). (n=8). B) Predicted IC50s of empty vector (grey), plasmid (pink) and chromosome integrated (dark blue) wild-type regulatory sequences. Significance shows results of pairwise comparisons of each variant to plasmid localised wild-type regulatory sequence (p-values adjusted using the Bonferroni correction for multiple comparisons, ***<0.001).

**Supplementary Figure S2:**
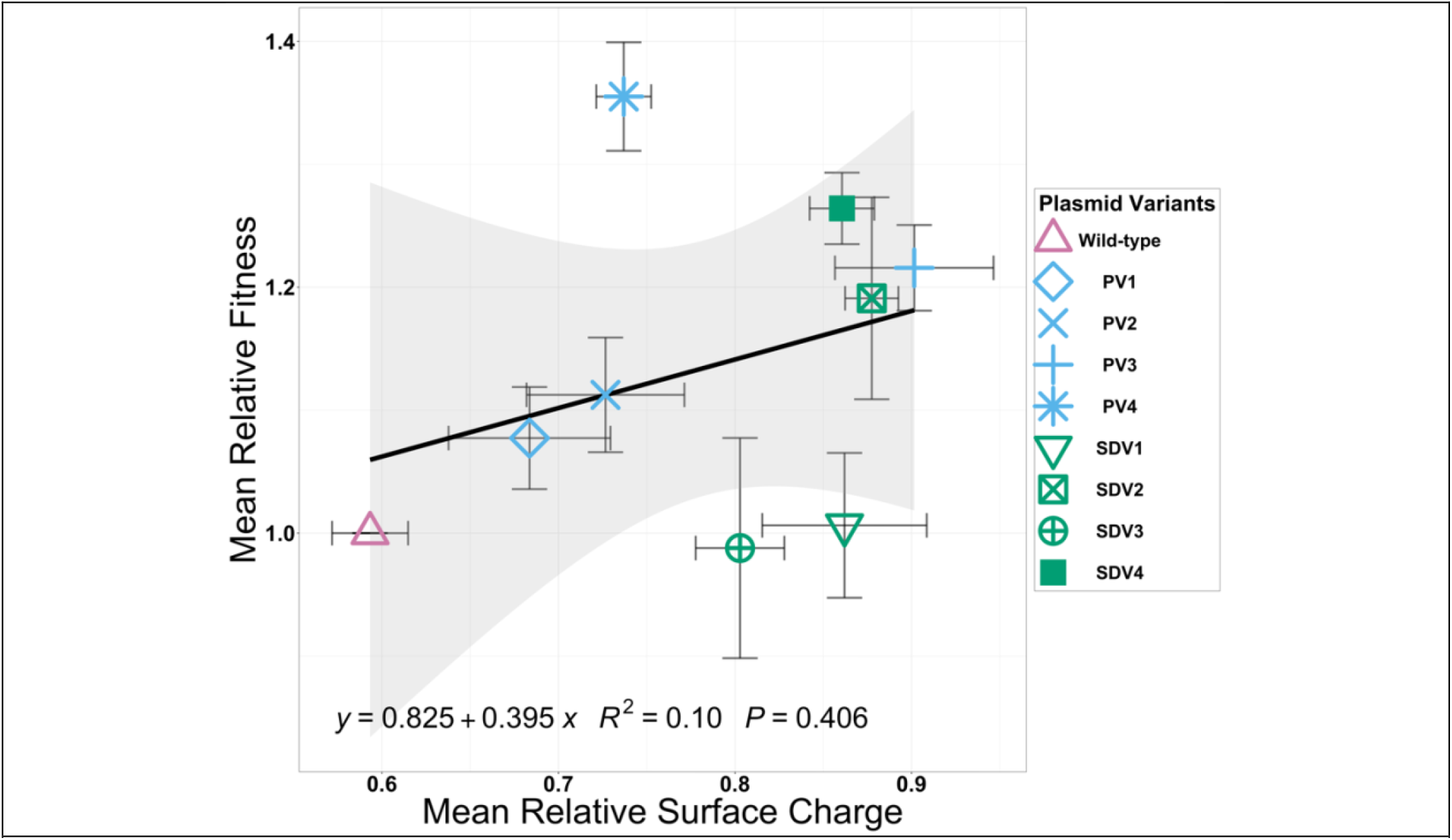
Scatterplot showing relationships between relative fitness (y-axis) and relative surface charge (x-axis) of regulatory variants and controls. Points assigned to variants/controls are shown in the legend (error bars = SE). Blank lines show linear regression model fitted to the data with confidence intervals in grey shading. Regression line equations, model r-squared and p-values indicated in black text.

**Supplementary Figure S3.**
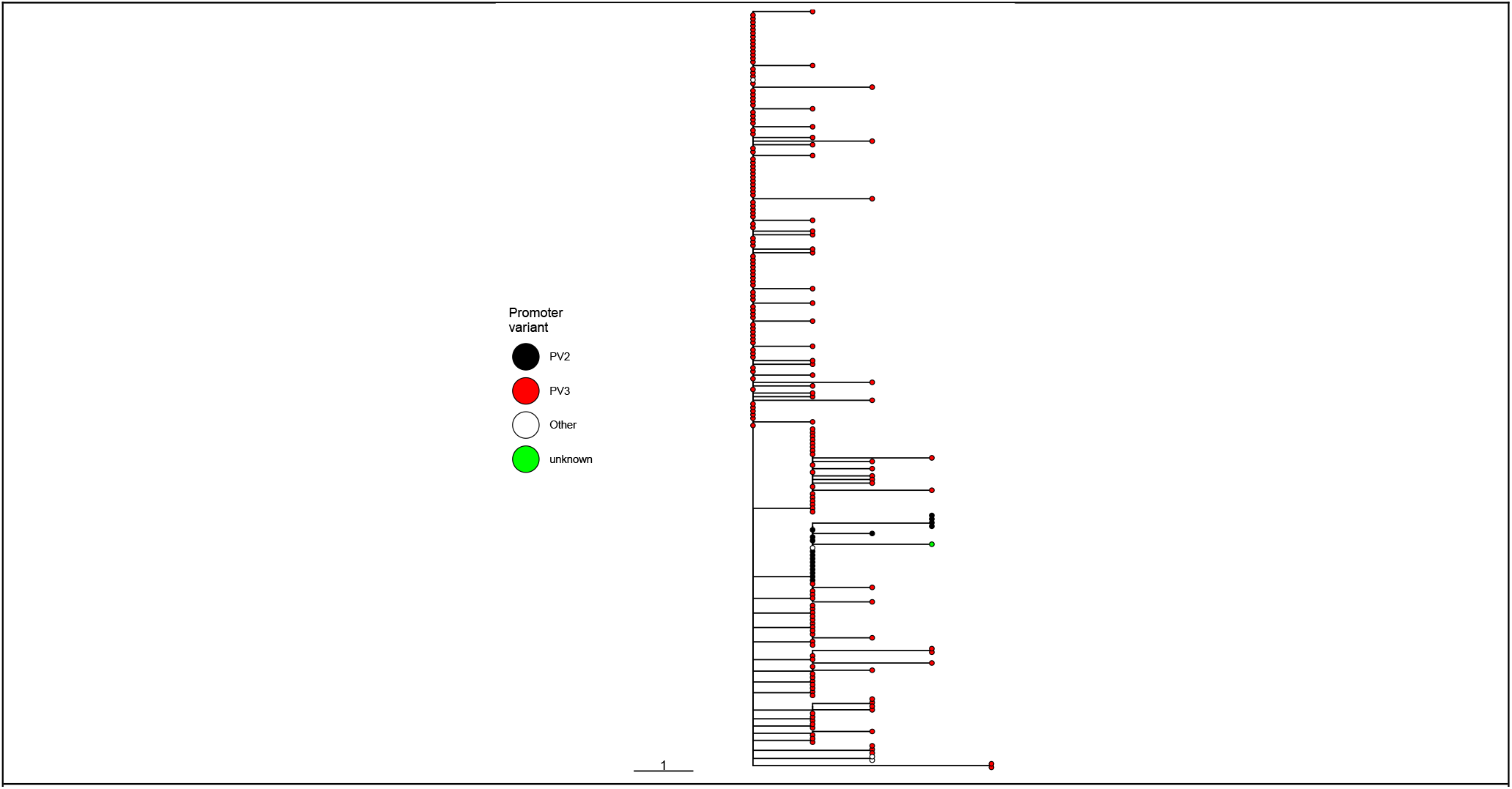
A phylogeny of IncX4 plasmids suggests two separate instances of fine-tuning acquisition. The tree is rooted to the reference plasmid used for the mapping (NCBI KU761327.1), with tip colour showing regulatory variant.

**Supplementary Figure S4:**
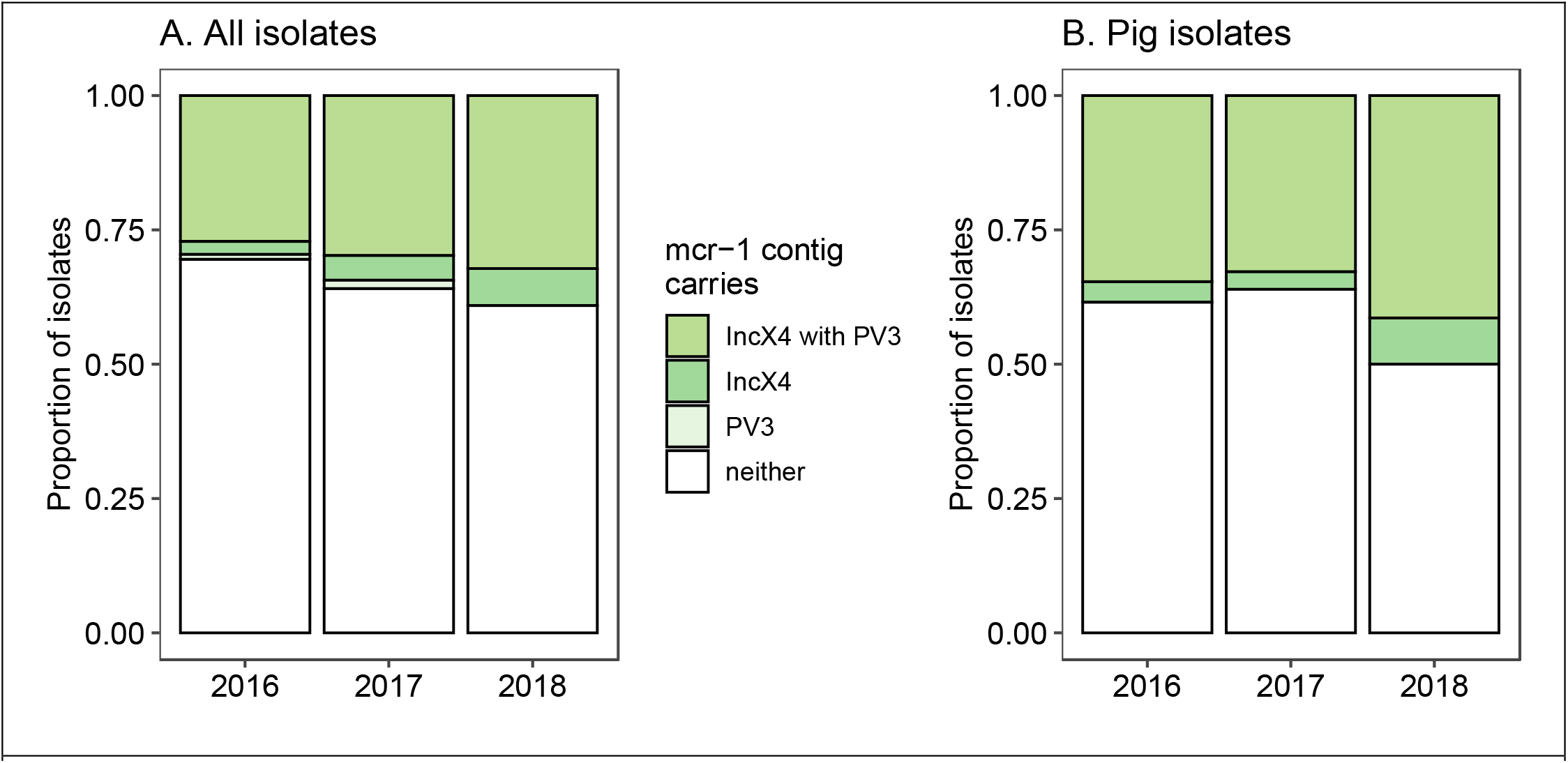
Increase in proportion of isolates carrying an IncX4 plasmid with PV3 from a genomic dataset of *mcr-1*+ isolates collected pre- and post-colistin ban. **A**) All isolates where the de novo assembly had an *mcr-1*-carrying contig (n=674). **B**) The subset of those isolates which were collected from pigs (n=197). Isolates are grouped by whether the *mcr-1*-carrying contig had a single IncX4 replicon and/or the PV3 regulatory mutation. Contigs with IncX4 and other replicons were not included within “IncX4”, because we are interested in the spread of plasmids similar to the reference plasmid which carries only IncX4.

**Supplementary Figure S5:**
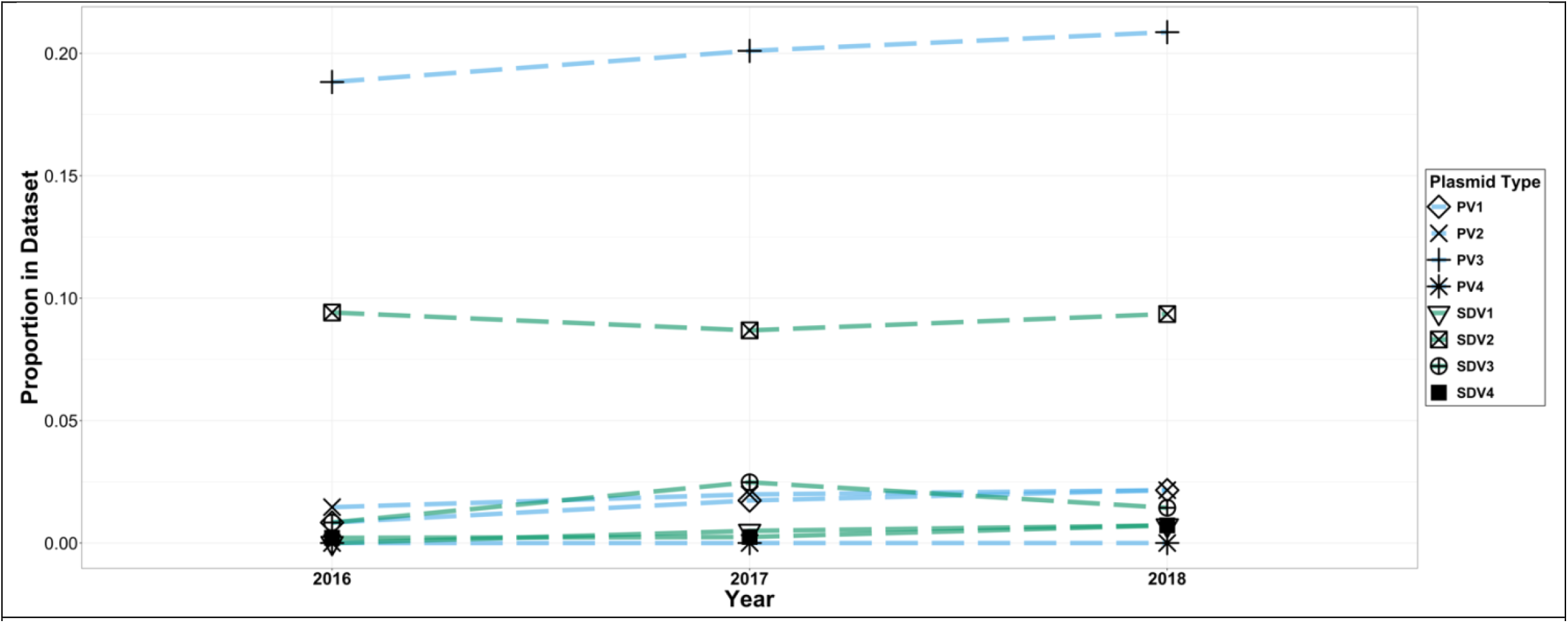
Proportion of promoter (blue) and SD (green) regulatory variant sequences in Shen dataset as separated by year.

## Statistical Tables

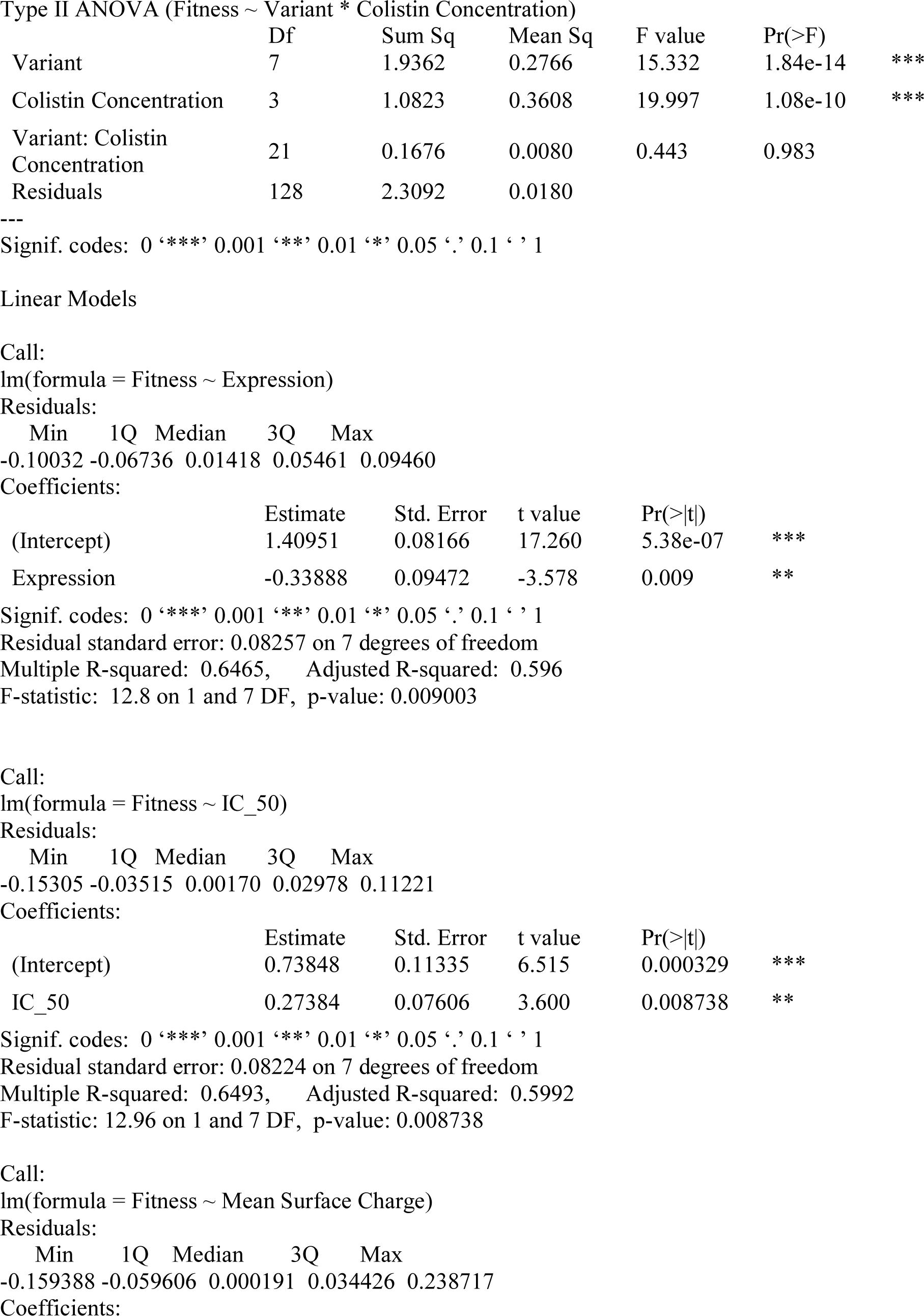
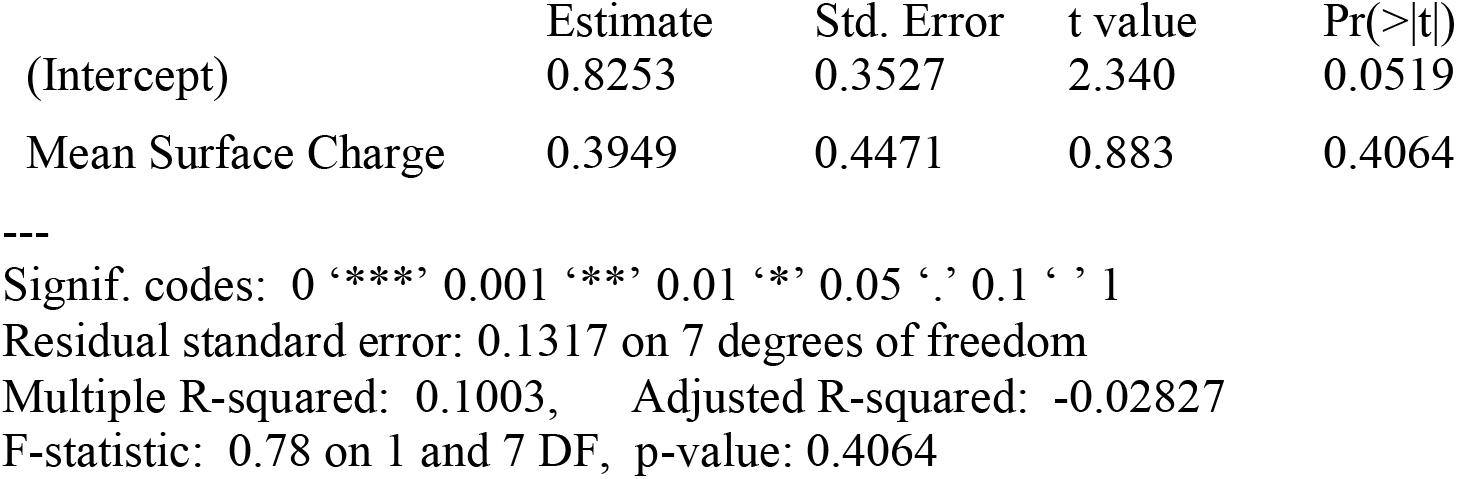

|  | Df | Sum Sq | Mean Sq | F value | Pr(>F) |  |
| --- | --- | --- | --- | --- | --- | --- |
| Variant | 7 | 1.9362 | 0.2766 | 15.332 | 1.84e-14 | *** |
| Colistin Concentration | 3 | 1.0823 | 0.3608 | 19.997 | 1.08e-10 | *** |
| Variant: Colistin Concentration | 21 | 0.1676 | 0.0080 | 0.443 | 0.983 |  |
| Residuals | 128 | 2.3092 | 0.0180 |  |  |  |

|  |  |  |  |  |
| --- | --- | --- | --- | --- |
| Min | 1Q | Median | 3Q | Max |
| -0.10032 | -0.06736 | 0.01418 | 0.05461 | 0.09460 |

Coefficients:
|  | Estimate | Std. Error | t value | Pr(>\|t\|) |  |
| --- | --- | --- | --- | --- | --- |
| (Intercept) | 1.40951 | 0.08166 | 17.260 | 5.38e-07 | *** |
| Expression | -0.33888 | 0.09472 | -3.578 | 0.009 | ** |

|  |  |  |  |  |
| --- | --- | --- | --- | --- |
| Min | 1Q | Median | 3Q | Max |
| -0.15305 | -0.03515 | 0.00170 | 0.02978 | 0.11221 |

Coefficients:
|  | Estimate | Std. Error | t value | Pr(>\|t\|) |  |
| --- | --- | --- | --- | --- | --- |
| (Intercept) | 0.73848 | 0.11335 | 6.515 | 0.000329 | *** |
| IC_50 | 0.27384 | 0.07606 | 3.600 | 0.008738 | ** |

|  |  |  |  |  |
| --- | --- | --- | --- | --- |
| Min | 1Q | Median | 3Q | Max |
| -0.159388 | -0.059606 | 0.000191 | 0.034426 | 0.238717 |

Coefficients:
|  | Estimate | Std. Error | t value | Pr(>\|t\|) |
| --- | --- | --- | --- | --- |
| (Intercept) | 0.8253 | 0.3527 | 2.340 | 0.0519 |
| Mean Surface Charge | 0.3949 | 0.4471 | 0.883 | 0.4064 |

Signif. codes: 0 ‘\*\*\*’ 0.001 ‘\*\*’ 0.01 ‘\*’ 0.05 ‘.’ 0.1 ‘ ’ 1
Residual standard error: 0.1317 on 7 degrees of freedom
Multiple R-squared: 0.1003, Adjusted R-squared: -0.02827
F-statistic: 0.78 on 1 and 7 DF, p-value: 0.4064

